# Non-homogenous axonal bouton distribution in whole-brain single cell neuronal networks

**DOI:** 10.1101/2023.08.07.552361

**Authors:** Penghao Qian, Linus Manubens-Gil, Shengdian Jiang, Hanchuan Peng

**Author notes:** These authors contributed equally (Penghao Qian, Linus Manubens-Gil).

## Abstract

We examined the distribution of pre-synaptic contacts in axons of mouse neurons and constructed whole-brain single-cell neuronal networks using an extensive dataset of 1891 fully reconstructed neurons. We found that bouton locations were not homogeneous throughout the axon and also among brain regions. As our algorithm was able to generate whole-brain single-cell connectivity matrices from full morphology reconstruction datasets, we further found that non-homogeneous bouton locations have a significant impact on network wiring, including degree distribution, triad census and community structure. By perturbing neuronal morphology, we further explored the link between anatomical details and network topology. In our in silico exploration, we found that dendritic and axonal tree span would have the greatest impact on network wiring, followed by synaptic contact deletion. Our results suggest that neuroanatomical details must be carefully addressed in studies of whole brain networks at the single cell level.

## Introduction

Neuronal morphology plays a fundamental role in determining the function of neuronal networks. Neurons are polarized cells conformed by the dendritic and axonal trees. Axons send connections to dendrites from other neurons through membrane specializations called synapses, which are structures that allow transmission of neuronal electrophysiological impulses. The architecture of the trees and the distribution of synapses within them determines the connectivity of neuronal networks. Changes in the structure of neurons can have a dramatic impact on cognition, and changes in dendrite shape and size have been associated with intellectual disability (Kulkarni and Firestein, 2012). However, understanding the effect of single neuron morphology on whole-brain circuit connectivity is still an open challenge.

Limited by resolution and time cost, current studies on the structural properties of whole-brain connectivity are mostly described from the mesoscale and macroscale perspective. Studies of human brain networks (Lynn and Bassett, 2019) using magnetic resonance imaging show there is a functional division in the brain at the anatomical level (Sherrington, 1907), called community structure (Hilgetag et al., 2000; Parente and Colosimo, 2020). This structure means that brain networks can be divided into sub-networks with specific cognitive functions (Azulay et al., 2016; Bassett et al., 2010; Lesicko et al., 2016; Sohn et al., 2011; Suárez et al., 2020; Taylor et al., 2017), with high node-density communities and sparse communities connecting them (Hilgetag et al., 2000; Sporns and Betzel, 2016; Sporns and Zwi, 2004; Sporns et al., 2000). Several experiments have also shown that the average path distance between nodes is much smaller in macroscopic brain networks than in random networks (Bullmore and Sporns, 2012; Liao et al., 2017; Van den Heuvel and Sporns, 2013), reflecting their small-world topology and the existence of central hubs (Gong et al., 2009; Sporns, 2022). This is thought to improve the segregation and integration of information within the brain (Deco et al., 2015), reducing the cost associated with information processing (Kaiser and Hilgetag, 2006; Latora and Marchiori, 2001). However, most of the experimental data obtained in these studies consists of ∼1mm sided voxels, which pool information from thousands of individual neurons.

At the mesoscale, the Allen Institute for Brain Science conducted a comprehensive study of the mouse brain connectivity, mapping the whole brain using population-based tracer injections (Oh et al., 2014). The study found that the clustering coefficients are close to those expected in a small-world network, while the degree distribution is close to a scale-free network (Oh et al., 2014). A refined connectivity analysis of the same experimental data showed global hubs in the mouse brain, including associative cortical areas, dorsal portions of the hippocampus and subregional portions of the basolateral and central amygdala (Coletta et al., 2020; Knox et al., 2018). The results also show highly connected central hub nodes interlinked with each other throughout the brain, supporting the efficient integration of otherwise segregated neural circuits. Neuromodulatory nuclei work as connector hubs and critical orchestrators of network communication at the fine granularity (Coletta et al., 2020). These studies have deepened our understanding of brain structure, but they still rely on measurements in large populations of tracer-injected neurons, missing the details of single neuron morphologies. Therefore, it is necessary to study neuronal morphology on the single-cell level as it is more meaningful in terms of finer and more essential brain network structures.

At the single-cell level, limited by the lack of experimental data, only a few studies have explored the impact of morphological details on whole-brain connectivity. In a study of the hippocampal trisynaptic circuit, a highly specialized topology has been found minimizing communication cost through information-processing hubs nested in a two-tier structure that manage the network traffic with strong resilience to random perturbations (Rees et al., 2016). However, A recent study of bouton-spine pairs in the rat barrel cortex found that most overlapping axons and dendrites were not connected (Udvary et al., 2022), indicating that the distribution of synaptic contacts in single neurons must be addressed in detail. However, it did not examine the impact of the detailed distribution of presynaptic contacts in full axon morphologies and it relies on the underlying assumption that pre-synaptic contacts are uniformly distributed. Complementing these results, a recent article has briefly explored the impact of perturbations on dendritic morphology on the wiring of the rat somatosensory minicolumn wiring. The study found that shortening or deleting the dendrites resulted in connection deficits in the neuronal network (Kanari et al., 2022), which is consistent with observations in neurological disease (Forrest et al., 2018). However, an exploration of how diverse perturbations in axonal trees and presynaptic contact distributions is missing, together with a graph-theoretical detailed analysis of the impact of perturbations on network topology.

The high-throughput generation of single neuron full morphology reconstructions in the mouse brain offers the possibility of exploring a novel approximation to single-cell whole-brain networks (Peng et al., 2021). Subsequent work has provided putative bouton locations throughout the fully reconstructed axonal trees (Jiang et al., 2022; Peng et al., 2023). The fact that neurons reconstructed from diverse brains are spatially registered (Qu et al., 2022) to a common coordinate framework (CCFv3) (Wang et al., 2020) allows us to analyze axon-dendrite potential connections at the whole-brain level with unprecedented anatomical detail. In fact, axonal boutons have been shown to be a proxy of structural neuronal connectivity (Grillo et al., 2013). However, reconstructing the full morphology of axonal projections and measuring the locations of single boutons in the context of the complete axonal tree was not possible until recently. For this reason, Peters’ rule (Peters and Feldman, 1976), a common approximation for quantifying neuronal connectivity at the cellular scale, has been used in the field of computational neuroscience. This rule assumes that there is a potential connection between a nearby axon and a dendrite, implying that synaptic connections are evenly distributed over the axonal and dendritic segments. However, it has been suggested that ground truth synaptic connectivity follows a nuanced Peter’s rule instead (Rees et al., 2017). From this perspective, the spatial distribution of pre-and post-synaptic sites and synaptic contact probabilities vary among diverse neuron types, finely tuning network connectivity.

We hypothesize that neuron morphology details determine network wiring. Specifically, we consider that distribution of axonal boutons throughout the axonal tree is not uniform, and such distribution determines network topology. Meanwhile, perturbation of specific morphological properties (i.e. neuron tree size, complexity, and density of axonal boutons) have a differential significant impact on network structure. To test this hypothesis, we devised an algorithm to generate single-cell networks in the whole brain using putative bouton locations and also simulated uniformly distributed boutons throughout the axon. First, we illustrate the biological relevance of axonal bouton distributions. With the networks we generated, we perform a detailed graph-theoretical analysis of the network structure and its dependence on axonal bouton distribution. To contextualize this information and explore its relevance in comparison to biologically plausible neural morphological alterations, we explored the impact of perturbing specific morphological features.

## Results

### Axonal bouton distribution is cell-type dependent

We analyzed putative axonal bouton locations obtained by another team (Peng et al., 2023) through automated detection of increased radius and intensity blobs in fully traced axons from neurons with somas located mainly in the Thalamic and Cortical regions. See Supplementary Table S1 for a summary of the number of neurons (N>20) in each analyzed CCFv3 brain region. The spatial distribution of the boutons is mainly determined by the axonal projection pattern of each cell type (Fig. 1A, B). To explore the biological relevance of the bouton distributions along the axons, we explored them in neurons with similar morphology and soma location in the brain (Fig. 1C top; Supplementary Figure 1A). To identify morphologically similar neurons, we used the Topological Morphology Descriptor (TMD), a method to encode the spatial structure of trees combining morphology and topology (Kanari et al., 2022; Li et al., 2017). The default TMD definition by Kanari et al. defines the barcode of a tree as the set of radial distances to the soma in the birth and death nodes of each branch of the tree. By measuring the distance between barcodes we obtained pairwise distances between all neurons in each brain region. To quantify the similarity in axon bouton distributions, we defined TMD bouton barcodes as the set of numbers of putative boutons enclosed within spheres with radii defined by the birth and death nodes of each branch of the tree. As expected, default TMD and bouton TMD distances are correlated, indicating that putative bouton locations are biologically meaningful (Figure 1C; Pearson correlation coefficient of 0.595 for default TMD vs. bouton TMD distances). Another observation that supports the validity of the data is that bouton density in somatosensory areas (obtained as the total length of axon divided by the number of boutons in these CCFv3 regions) is 0.0778 boutons per μm which is close to 0.061 boutons per μm experimentally measured in adult mice using serial section electron microscopy (SSEM) data (Grillo et al., 2013).

**Figure 1.**
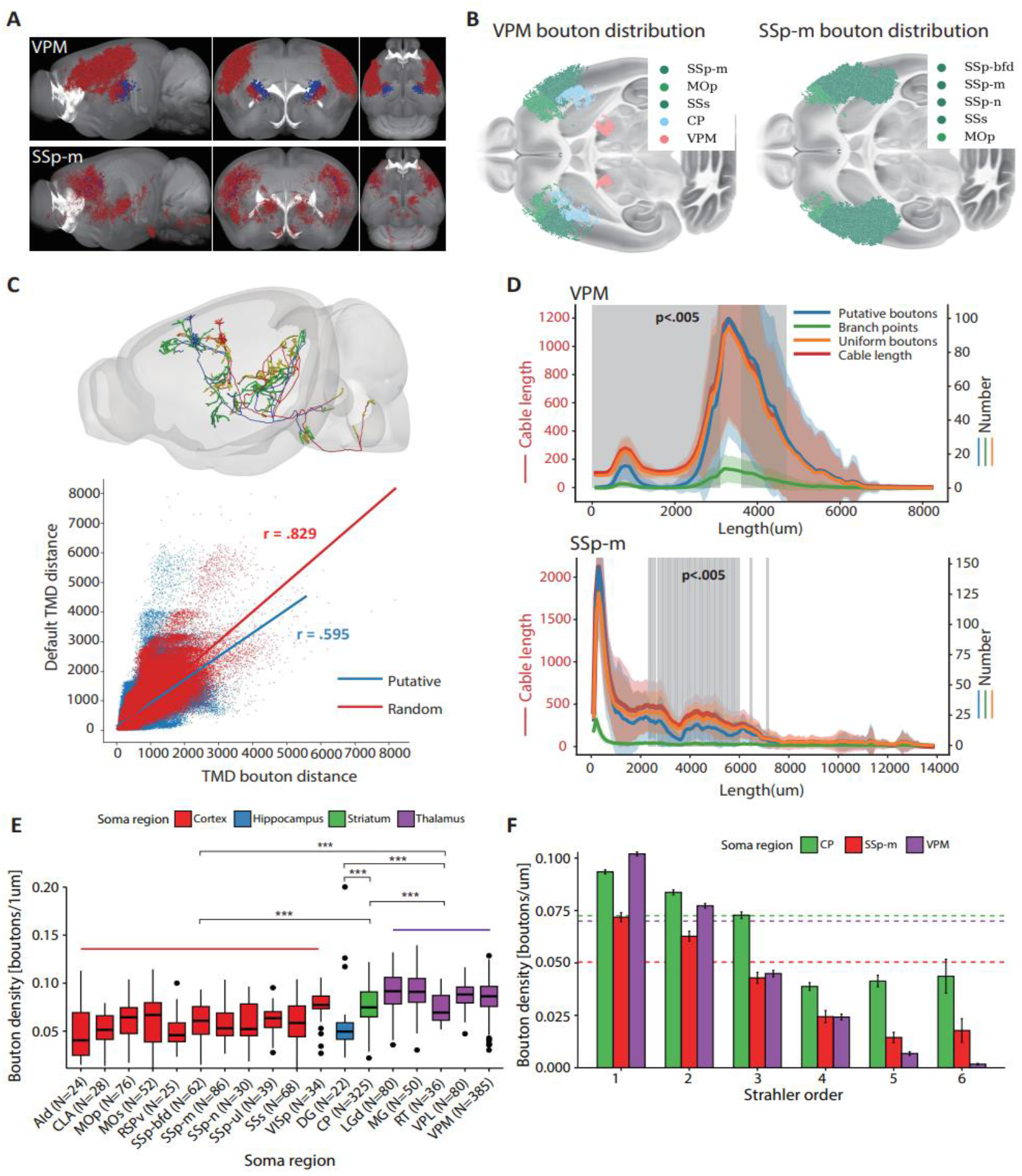
Axonal bouton distribution is cell-type dependent. (A) 2D sagittal, coronal, and horizontal projections of putative bouton locations (in red) from two sets of neurons with somas (in blue) located in the Ventral posteromedial nucleus of the Thalamus (VPM) and mouth Primary somatosensory area of the cortex (SSp-m) based on the CCFv3 parcellation (N_VPM_=379, N_SSp-m_=78). See Supplementary Table S1 for a complete list of acronyms. **(B)** Horizontal projection of bouton locations in the top five regions most innervated by VPM and SSp-m neurons. Cortical regions are colored in shades of green, caudoputamen (CP) in the Striatum, in blue, and VPM in the Thalamus in pink. **(C)** Top: morphologically similar neurons have analogous bouton distributions throughout the axons. Bottom: Scatterplot of the Topological Morphology Descriptor (TMD) distances between pairs of neurons with somas in the same region using default TMD (x axis) or TMD bouton (y axis; see methods). The plot shows pairwise distances for neurons with putative bouton locations (blue) and analogous measures for uniformly distributed boutons (red). “r” values specify Pearson correlation coefficients. **(D)** Sholl analysis of the number of boutons, cable length and number of boutons for neurons with their soma in VPM (top) or SSp-m (bottom). Statistically significant differences between uniform and observed bouton distributions are indicated with a grey shadow; paired-samples t-tests random vs. observed bouton number p<0.005. **(E)** Barplot of the fitted average bouton density (see Methods) for all neurons in each of the brain region with more than 20 neurons. Pairwise t-tests between brain areas; *** p<0.001 **(F)** Barplot of bouton densities at different Strahler orders for neurons with somas in CP, SSp-m, and VPM. Bars indicate mean ±s.e.m. Dashed lines indicate the average bouton density throughout the full axonal tree.

To explore whether bouton distributions are non-uniform we defined null uniform distributions of bouton locations for each neuron (see methods). Interestingly, the null distribution showed a higher correlation with default TMD than the measured distribution of putative boutons (Fig. 1C bottom and Supplementary Fig. 1A; Pearson correlation coefficients of 0.829 vs. 0.595 for random and measured respectively; one-way ANCOVA F(1,3109929)=0.022, p=7.17e-9). This is explained by the fact that our bouton TMD definition is highly sensitive to differences in the total numbers of boutons of morphologically similar neurons (see Supplementary Figure 1B). Correlations between TMD distances segregated for neurons with their somas in specific brain regions show analogous results (Supplementary Figure 1C). A Sholl analysis of axon length, branch points and number of boutons showed (see methods) putative bouton distributions significantly differ from an uniform distribution (Fig. 1D; paired-samples t-tests random vs. putative bouton number p<0.005). Specifically, the boutons of neurons in the thalamus (VPM) are preferentially located at the distal axon and have low bouton density near the soma. Conversely, neurons in the somatosensory cortex (SSp-m) show a distribution of boutons that is close to uniform for both proximal and distal axonal branches, and significantly lower than uniform in the middle section (see other cell types in Supplementary Figure 2A and 2B).

**Figure 2.**
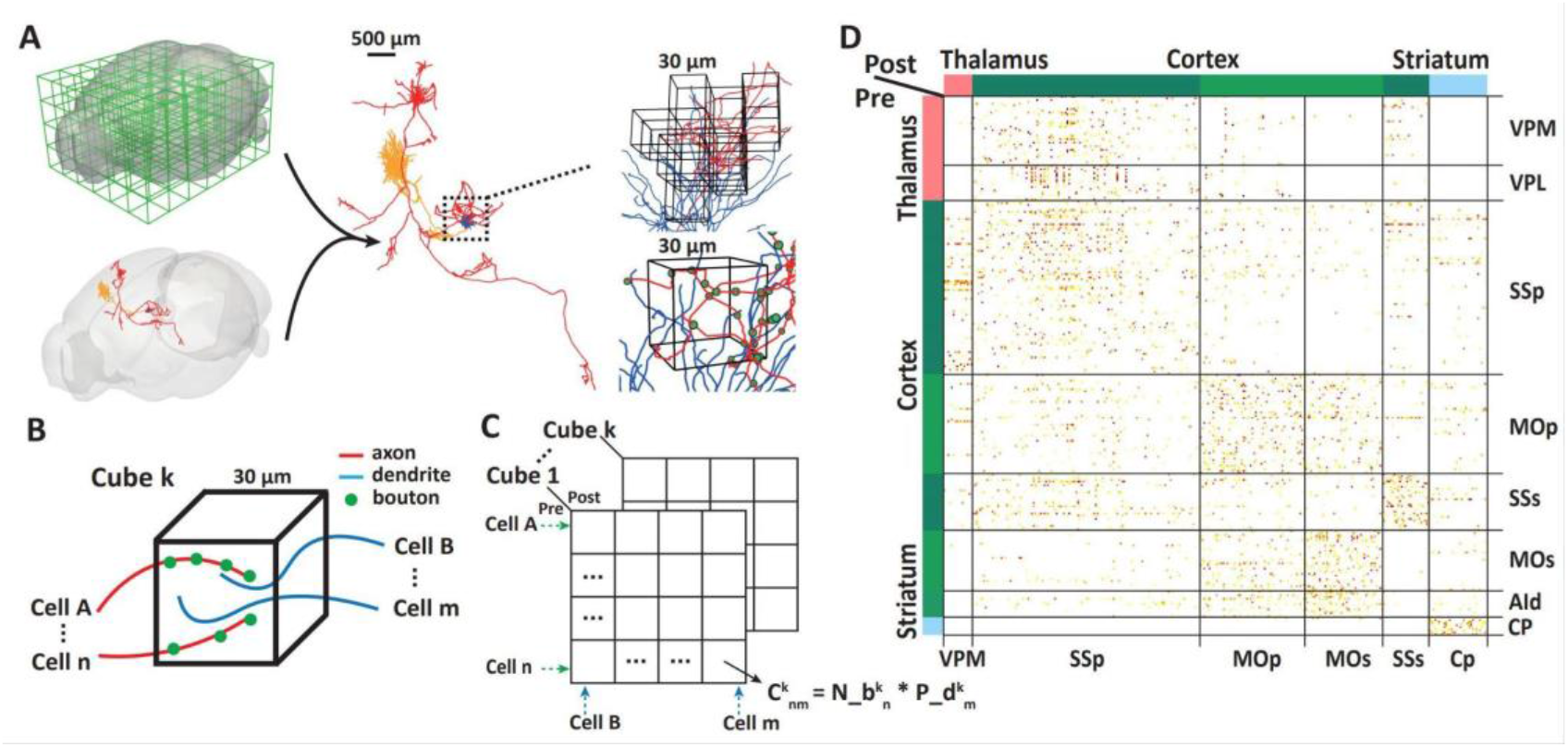
Generation of connectivity matrices based on full neuron reconstructions. (A) 3D rendering of the division of the whole brain into cubes of 30 μm units (top left) and full neuron reconstruction registered to CCFv3 (bottom left). Close-up rendering of a pair of neurons (dendrites in blue and axons in red and orange; middle). Rendering of a region of interest where axons and dendrites are close-by in CCFv3 space (top right) where the length of dendrite and The number of axonal boutons (green dots) can be obtained within each cube (bottom right). **(B)** Schematic visualization of the co-existence of axon and dendrite within each cube, which define connectivity together with axonal boutons. **(C)** Schematic visualization of our procedure to obtain the connection strength between each pair of neurons in a cube according to the number of boutons and the dendrite length of each post-synaptic neuron found in the cube. **(D)** Visualization of a subset (304 and 306 pre-and post-synaptic neurons respectively) of the whole brain single-cell connectivity matrix.

Obtaining small fragments of axon and quantifying axonal bouton density is a common practice in electron microscopy studies (Casas-Torremocha et al., 2019; Grillo et al., 2013; Rodriguez-Moreno et al., 2020). Our results indicate that those estimates may vary among distal and proximal axonal trees. To provide improved estimates based on our detailed data, we fitted an uniformly distributed bouton curve to the observed distribution of putative boutons (see Methods). Neurons with their somas in different brain regions showed different average bouton density values (Fig. 1E; paired samples t-tests Cortex vs. Thalamus p<2e-16, Cortex vs. Striatum p<2e-16, Cortex vs. Hippocampus p=0.98, Hippocampus vs. Thalamus p=1.1e-9, Hippocampus vs. Striatum p=9.2e-5, Striatum vs. Thalamus p=3.9e-11). The average bouton density of the analyzed cell types ranged from 0.029 boutons per μm to 0.104 boutons per μm.

As previously described for neocortical inhibitory neurons (Budd et al., 2010), we found that lower Strahler order (Nebel, 2000; Strahler, 1957) segments (tip or close-to-tip segments) have the highest density of boutons, decreasing towards higher Strahler order segments, especially above 2 (Fig. 1F). Interestingly, the Strahler order distribution of bouton densities varied among neurons with somas in different brain regions (Fig. 1F; all pairwise t-tests p<0.05 except for SSp-m vs. VPM in Strahler orders 3 and 4). The low variance in the bouton density within each Strahler order indicates that those can be considered homogeneous.

To investigate the details of single-cell morphology, including the size and complexity of axons and dendrites, and the specific effects of the number and distribution of boutons on the network, we devised a method for constructing single-cell networks based on full neuron tracings. We argue that if there are axon boutons and dendrites in close spatial proximity, then they have a high probability of producing synaptic contact (Rees et al., 2017). To generate connectivity matrices, we divided the whole brain into 30*30 μm cubes and measured the axon length, dendrite length, and bouton number of each neuron within each cube (Figure 2A). We consider that when both axons and dendrites are present in a cube containing boutons, there is a potential connection (Figure 2B). Then we defined the connection strength (Figure 2C) as follows. Considering that an axon may connect to many dendrites in the same cubic volume, we set the strength of the connection proportional to the number of boutons and to the proportion of the length of each dendritic tree. Specifically, the connection strength of the presynaptic neuron *n* with the postsynaptic neuron *m* in a single cube is defined as:

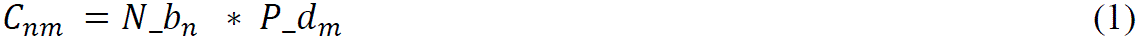

Where the number of boutons of neuron *n* is *N*_*b*_*n*_ and the proportion of dendrite length of neuron *m* is *p*_*d*_*m*_. We obtained the full brain single-cell connectivity matrix by adding the results obtained on all cubes covering the mouse brain in CCFv3 (Figure 2D).

We applied this network generation method to our single-cell data with observed bouton locations (***observed* network**). To explore the relevance of the bouton distribution on the network topology, we also generated an ***uniform* network** setting boutons throughout the axon using the average bouton density for each cell type we obtained previously (Figure 1E).

A comparison between our *observed* connectivity matrix and the mesoscale connectome obtained previously with tracer injections (Oh et al., 2014) shows similar connectivity clusters (Supplementary Figure 3A), suggesting that even though our single neuron data accounts for a very small percentage of the network, it can recapitulate the mesoscopic structure. The *uniform* network shows a bias towards increased connectivity, with 1847 nodes and 22882 edges compared to the *observed* network with 1781 nodes and 14056 edges. The small difference in the number of nodes is due to the removal of neurons with 0 potential connections. Circular plots of the network structure show that there is a large number of potential connections between the cortex, thalamus, and striatum, as well as local potential connectivity (Figure 3A).

**Figure 3.**
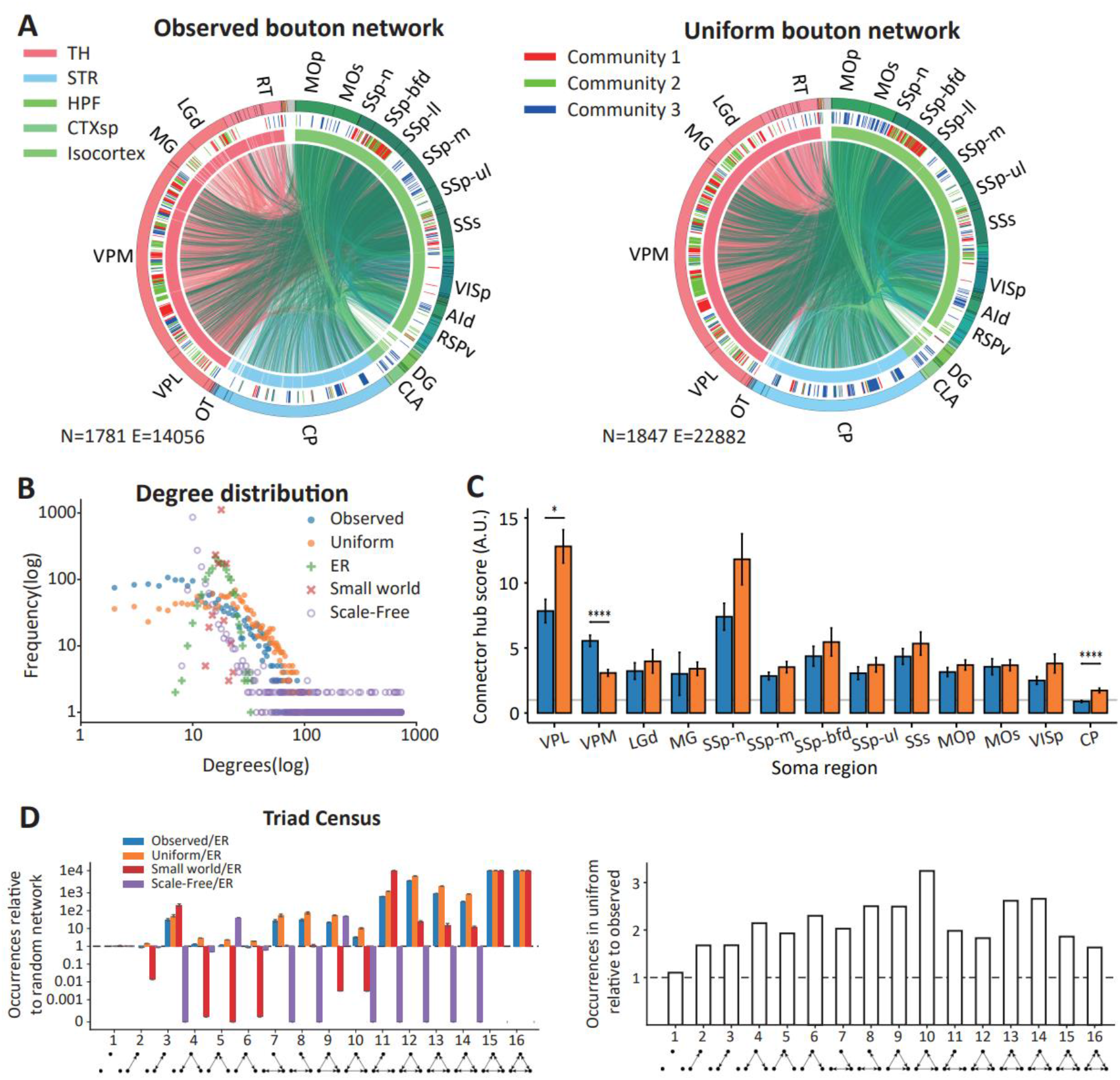
Bouton distribution tunes network topology and influences community structure. (A) Circular plot visualization of the single-cell networks constructed based on *observed* (left) and *uniform* (right) bouton data. We assigned the colors in the outer circle by the soma location of each neuron in the network. The middle circle indicates the largest three communities obtained using the Leiden algorithm (Traag et al., 2019), and the inner circle indicates broad brain regions Thalamus (TH), Striatum (STR), hippocampal formation (HPF), cortical subplate (CTXsp), and Isocortex. The lines crossing the center of the circle indicate potential connections between individual neurons. We colored them according to the soma location of the pre-synaptic neuron. **(B)** Scatter plot of the degree distributions of *observed* (blue), *uniform* (orange), Erdös-Rényi (ER; green), small-world (red), and scale-free (purple). (**C**) Barplot of connector hub scores for the soma regions with at least 50 neurons for *observed* (blue) and *uniform* (orange) networks. The gray line indicates connector hub score 1, which indicates the same proportion of edges within and across regions. Pairwise t-tests * p<0.05, **** p<0.0001. (**D**) Barplot of triad occurrence is relative to ER networks with the same number of nodes and edges (left) of *observed* (blue), *uniform* (orange), small-world (red), and scale-free (purple) networks. Bars indicate mean values ± standard deviation. Barplot of triad occurrence in the *uniform* network relative to *observed* (right). The dashed lines indicate no difference (ratio=1). The bottom row shows schematic visualizations of the network motifs identified with each index from simple to complex.

To explore the community structure, we used the Leiden community detection algorithm (Traag et al., 2019) (see methods) on the two networks. Remarkably, the circular plots show that the bouton distribution tunes the community structure in the network. In particular, we assessed the number of neurons in each community and found that, in the *observed* network, the largest community of neurons (N=206) belongs to the thalamo-cortical loop in the left hemisphere (mainly from VPM to SSp-bfd in cortex; Figure 3A and Supplementary Figure 4A). However, this loop is under-represented in proportion in the *uniform* network, being the third largest community (N=215) (Supplementary Figure 4B). The second largest community in the *observed* network is defined by the right hemisphere thalamo-cortical loop with a higher representation of VPL and MG in the thalamus and other cortical areas such as SSs and RSPv (N=179). The most similar community in the *uniform* network is larger in proportion (N=256), having an over-representation of VISp neurons. The third largest community in the *observed* network is the cortex-thalamus-striatum loop of cortical neurons with CP and VPM (N=172). This community is the second largest in the *uniform* network (N=234), where motor cortex, CP and AId areas are over-represented. Conversely, for this community, SSp neurons are under-represented in the *uniform* network.

Previous studies show that there are some simple organizational patterns such as the degree distributions (Barabási and Albert, 1999) in brain networks that can be explained using network models (Lynn and Bassett, 2019). To explore this, we generated three artificial networks: random, small-world, and scale-free with the same number of nodes and edges as the *observed* network for comparison. Given that Oh et al. (Oh et al., 2014) showed that the mouse brain network has Small-World topology traits (namely large clustering coefficienc and short average path length (Watts and Strogatz, 1998)), we tested whether the *observed* and *uniform* networks follow this trend. In the random network, the average path length is 3.852 and the clustering coefficient is 8.96e-3, compared to the *observed* average path length of 4.828 (*uniform*: 4.257) and clustering coefficient of 0.184 (*uniform*: 0.195) in the *observed* network (Supplementary Table S4). These resultsis are consistent with previous data ((Oh et al., 2014)). However, the model that best approximates the degree distribution of the *observed* and *uniform* networks is the scale-free network (Figure 3B; Spearman correlation *observed* vs. scale-free r(1779)=0.242, p<0.001; *observed* vs. random network r(1779)=0.029, p>.1; *observed* vs. small-world network r(1779)=0.0136, p>0.1). However, degree distributions strongly differ among all cases (see Supplementary Table S3 for pairwise two-sample Komogorov-Smirnov tests; p<1e-83). When comparing *observed* with *uniform*, the degree distribution shows an increased proportion of high-degree nodes (two sample Kolmogorov-Smirnov D(1779)=0.297, p<.001). To better understand this difference, we tested whether high-degree nodes are different among the two networks. We obtained hub and authority scores for anatomically defined brain regions (Kleinberg, 1999). The brain regions with the highest hub scores are SSp-m and VPM in both networks (Supplementary Table S4). Authority nodes are mainly cortical regions (including SSp-n, SSs, SSp-un, SSp-m, and SSp-bdf) for the *observed* network, but also include CP, VPL, and VPM in the *uniform* network. Those can be linked to the integration and segregation of the networks (Lord et al., 2017) by measuring the ratio between local and inter-region potential connections, which identify provincial and connector hubs. We found that the *uniform* network tends to have higher connector hub scores except for VPM (Figure 3C; pairwise t-test *observed* vs. *uniform* VPL p=0.017, VPM p=8.1e-7, CP p=1.3e-5). This also suggests that the specific distribution of axonal boutons can have functional implications, and is relevant to understand the contribution of each brain region in information transmission and processing throughout the brain.

The pattern of potential connections between triads is the most basic motif forming the networks (Holland and Leinhardt, 1977), and its distribution reflects the rules of neuronal connectivity (Udvary et al., 2022). We found that *observed* and *uniform* networks have similar triad distributions (Supplementary Figure 4C; two-sample Kolmogorov-Smirnov D(14)=0.3125, p=0.42), and both strongly differ from random, small-world and scale-free networks (Figure 3D top and Supplementary Table S4), showing more prominent feedback and complex potential connections compared to the expected occurrence in random networks (Figure 3D and Supplementary Table S4). This result is consistent with previous observation in cortical microcircuits ((Udvary et al., 2022)). However, the *uniform* network overestimates all network motifs in comparison to *observed*, with an average ratio of 2.1 (Figure 3D bottom), highlighting a methodological artifact in studies assuming homogeneous axonal bouton distributions.

Overall, our results suggest that the detailed distribution of axonal boutons is a relevant determinant of the network topology. Taking into account that the basic function of brain networks is to transfer and store information, we measured cost (defined as the total number of boutons in the network), routing efficiency (Goñi et al., 2014), and storage capacity (Poirazi and Mel, 2001) which are topological correlates of the network functional performance (Avena-Koenigsberger et al., 2014) (see methods). Since the *uniform* network has more edges and stronger potential connections, its cost, routing efficiency, and storage capacity are 1.563, 1.561, and 1.595 times higher than the *observed* network respectively (Supplementary Figure 4D). But the average routing efficiency and storage capacity per bouton, which is the value divided by cost and is representative of the Pareto optimality of the network (Avena-Koenigsberger et al., 2014), did not change (normalized routing efficiency and storage capacity are 9.6e-3 A.U. and 0.38 A.U. respectively for both networks). Thus, it remains an open question whether non-random distributions of axonal boutons imply any functional advantage from an energy optimization perspective.

To analyze the impact of putative morphological alterations relevant to cognitive impairments (Baloyannis, 2009; Emoto, 2011; Huang and Rasband, 2018; Koleske, 2013; Kweon et al., 2017; O’Keeffe and Sullivan, 2018), we perturbed the networks as follows: scaling neuron size, pruning neuron branches, and removing axonal boutons (see methods). Tree span has the greatest impact on the network, implying a marked shift towards lower degree potential connections (Figure 4C left, Supplementary Figure 5A; Kolmogorov-Smirnov unperturbed vs. scale all 0.5 D(86)=0.477, p=6.5e-9) and orders of magnitude lower occurrence of complex network motifs (Figure 4C right; Kolmogorov-Smirnov unperturbed vs. scale all 0.5 D(14)=.438, p=.0933). Conversely, pruning and bouton deletion, even when reducing the number of branches or boutons to half, had a modest impact on degree distribution (Figure 4C left; Kolmogorov-Smirnov unperturbed vs. prune all 0.5 D(86)=0.198, p=0.07; Kolmogorov-Smirnov unperturbed vs. bouton delete all 0.5 D(86)=0.093, p=0.85) and the triad census (Figure 4C right; Kolmogorov-Smirnov unperturbed vs. prune all 0.5 D(14)=.25, p=.716; Kolmogorov-Smirnov unperturbed vs. bouton delete all 0.5 D(14)=.125, p=.999).

**Figure 4.**
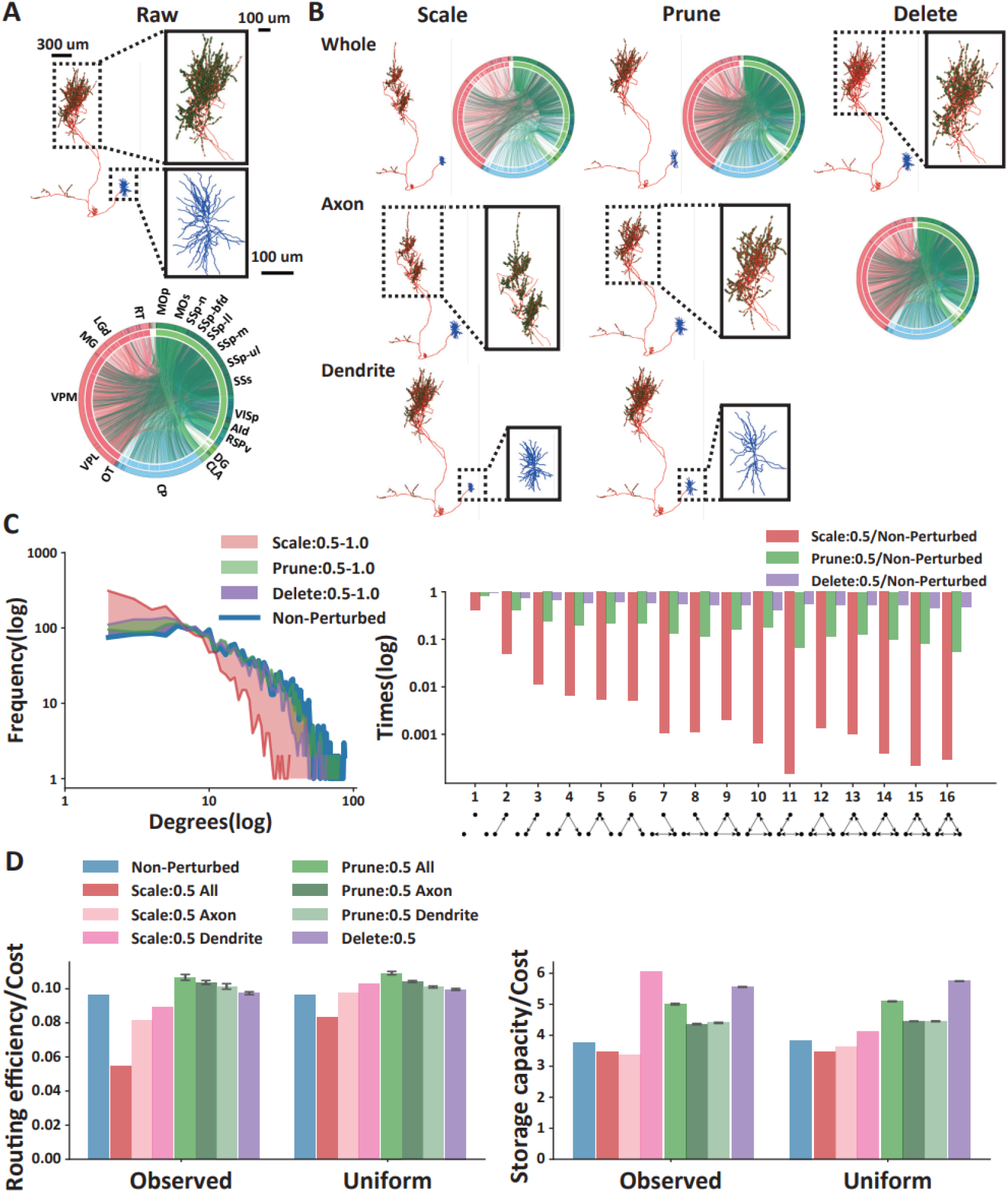
Dendritic and axonal tree spans are the main determinants of network topology. (A) Representative example of a full morphology neuron (top) with magnifications of dendritic (blue) and axonal (red) arbors. The axonal arbor magnification shows axonal boutons as green dots. The circular plot of the *observed* unperturbed network (bottom). Colors in the outer circle indicate the soma location of each neuron. The inner circle indicates broad brain regions Thalamus (TH), Striatum (STR), hippocampal formation (HPF), cortical subplate (CTXsp), and Isocortex. The lines crossing the center of the circle indicate potential connections between individual neurons, colored according to the soma location of the pre-synaptic neuron. **(B)** Representative visualizations of single neuron perturbations in both dendritic and axonal arbors of the neurons (top row), only in axonal arbors (middle row), and only in dendritic arbors (bottom row). The perturbations include scaling of the tree span (left row), pruning of branches (middle column), and deletion of boutons (right column). Scale bars=100 um. **(C)** Line plot of the degree distributions (top) for the unperturbed *observed* network (blue) and for scale (orange), prune (green), and bouton deletion (purple) perturbations in both dendritic and axonal arbors. The colored shadows indicate the range in which degree distributions vary with each perturbation with ratios between 0.5 and 1. Bar plot (bottom) of triad occurrence relative to the *observed* unperturbed network for scale (orange), prune (green), and bouton deletion (purple) perturbations. The bottom row shows schematic visualizations of the network motifs identified with each index from simple to complex. **(D)** Bar plots of routing efficiency (left) and storage capacity (right) divided by the cost (number of potential contacts) for the unperturbed *observed* and *uniform* networks (blue) and after scale (red), prune (green), and bouton deletion (purple) perturbations. Bars indicate mean values ±standard deviation.

Similarly, we investigated the community structure of different networks after perturbation. Pruning of axonal and dendritic branches, or bouton removal does not imply marked differences in the community structure (Supplementary Figure 6A, 6B, and 6C). The top two communities are still the cortico-thalamic loop of both hemispheres and the third is a cortico-thalamic-striatal loop. However, the scaling of dendritic and axonal trees sharply reduced the number of potential connections (having 3955 edges compared to 14056 in the *observed* network), strongly impacting the community structure (Supplementary Figure 6D). While dendritic scaling strongly reduced the number of potential connections (average of 373 potential connections per community) in comparison to axonal scaling (average of 760 potential connections per community), the change in the community structure for axonal scaling implied losing the thalamic connection in the thalamo-cortical-striatal circuit (Supplementary Figure 6C).

When we measured routing efficiency in the networks perturbed with pruning and bouton deletion, we found that the reduction in routing efficiency provoked by the perturbations is compensated by the reduction in cost (Figure 4D left, absolute value in Supplementary Figure 5B). Interestingly, the impact of both axonal and dendritic tree downscaling implied a marked reduction (43.35%) of the routing efficiency per unit of cost in the *observed* network, but had only a subtle impact (13.74%) in the *uniform* network (Figure 4D left, Supplementary Figure 5C). This indicates that the impact of tree span perturbations may be underestimated in generative models not taking into account precise pre-synaptic connection distributions in the axon. Morphological perturbations showed a subtle increase in storage capacity per unit of cost (Figure 4D right, Supplementary Figure 5C). This is explained by the fact that the approximation we used for storage capacity is mainly dependent on the combinatorics of diverse afferents on the same post-synaptic neurons (Poirazi and Mel, 2001). Interestingly, the *observed* network showed a storage capacity per unit of cost markedly higher than in the *uniform* network (60.99%) when only dendritic trees are downscaled (Figure 4D right). This is the only case in which the observed distribution of putative boutons appears to have a strong impact on network topology. This result suggests that the synaptic targeting of diverse axonal arbors is precisely matched with postsynaptic dendrites, implying that even when strongly reducing dendritic tree size, post-synaptic neurons can still keep receiving inputs from a diverse set of incoming axons.

To assess the robustness of our results taking into account that the experimental data is highly sparse, we removed half of the neurons in the *observed* network (Supplementary Figure 7A). We found that both the degree distribution and triad census did not vary in comparison to the *observed* network (Supplementary Figure 7B and 7D), while the community structure and routing efficiency and storage capacity per unit of cost markedly increased (by a factor of 2.02 and 1.71 respectively; Supplementary Figure 7C). This suggests that increasing the numbers of neurons used to generate whole-brain connectivity networks is necessary to accurately assess circuit architecture.

## Discussion

Quantifying the precise distribution of pre-synaptic contacts in full axons of mammal neurons has been addressed in some studies that require arduous manual effort for their annotation (Casas-Torremocha et al., 2019; Grillo et al., 2013; Rodriguez-Moreno et al., 2018, 2020). In this work, we leverage automatically identified putative axonal bouton locations obtained by our team (Peng et al., 2023). In accordance with previous studies, our analysis shows that pre-synaptic contact locations are not homogeneous throughout the axon (Brown et al., 2012; De Paola et al., 2006) and that they vary among brain regions (Karube et al., 2004; Rodriguez-Moreno et al., 2020). The consistency we found in morphologically similar neurons suggests that the putative bouton connection data we used is sufficient as a first approximation. Similarly, the low variance we found in our quantification of increased axonal bouton density in terminal branches is supported by previous evidence in local axonal trees, which has been suggested to enhance temporal economy and precision in neocortical inhibitory axonal trees (Budd et al., 2010). The fact that this phenomenon is found for all the neurons we analyzed, indicates it is a fundamental principle determining pre-synaptic contact distribution. Still, further refining the methods to take into account multiple synapses in single boutons and pre-synaptic sites in the absence of boutons will be necessary in the future. It is worth noting that the aim of this work is not to generate accurate and complete connectivity matrices but to explore the relevance of non-homogeneous pre-synaptic contact distributions on the network structure.

The fact that simple dendrite-axon colocalization and averaged synapse densities obtained from Electron Microscopy experiments are not sufficient to approximate network connectivity has already been addressed by previous publications (Rees et al., 2017). One relevant aspect of our work is its special focus on long-range projections, together with the development of the method to generate whole-brain single cell connectivity matrices. Our algorithm for generation of connectivity matrices based on full morphology neuronal reconstructions is open-source and our scripts conform to a pipeline to generate full-brain networks with datasets that are expected to grow exponentially in the future. Our comparison between *observed* and *uniform* bouton distributions and their impact on network structure supports the claim that Peters’ rule is an over-simplified model, which can not truly reflect the differences in connections between brain regions and between neurons. It overestimates the possibility of the existence of connections, which leads to an overall bias in the properties of the network and significant differences in its community structure and provincial vs. connector hub score, which is relevant for functional integration and segregation (Lynn and Bassett, 2019). These results are consistent with previous findings in the cortical network architecture, where most overlapping axons and dendrites are not connected, and the more dendrites from different neurons the axon is exposed to, the less probability of connection exists (Udvary et al., 2022).

The network properties we found in single-cell networks also complement our understanding of neuronal wiring rules. The *observed* bouton network we analyzed shows increased occurrence of feedback, and complex network motifs than what would be expected in random networks, which is consistent with the result found in the barrel cortex (Udvary et al., 2022). Our results indicate that such a pattern in the distribution of network motifs is not unique to cortical networks but is also present in the thalamo-cortical loop. Also in this study, the network was found to have small-world properties, which was also confirmed in our single-cell network. The degree distribution is close to scale-free and the clustering coefficient is small (Oh et al., 2014).

Existing studies suggest that neurological diseases such as intellectual disability, autism spectrum disorder, epilepsy, schizophrenia, and bipolar disorder, are accompanied by a decrease in dendrite branches and spines with atrophy of the dendrite morphology (Baloyannis, 2009; Emoto, 2011; Koleske, 2013; Kulkarni and Firestein, 2012; Kweon et al., 2017). The most influential of these changes is the decrease in the spine and the change in morphology (Forrest et al., 2018). Deformation and damage of axons can also lead to various neurological diseases (Huang and Rasband, 2018; O’Keeffe and Sullivan, 2018). This phenomenon can be seen in our perturbations. The biggest impact on the network properties is the dendritic and axonal tree span scaling. And the studied network is more robust to bouton removal and pruning of branches, which is also supported by previous literature (Aerts et al., 2016). All operations on dendrites have a greater impact than on axons for degree and network motif distributions. However, it is interesting to note that dendrite downscaling shows increased storage capacity per number of connections for the *observed* network, while the same quantity does not change for the *uniform* network. This is an unexpected result suggesting that the precise location of axonal boutons in diverse axonal arbors allows to keep high input combinatorics in single dendritic trees even with marked downscaling of the dendritic span.

### Limitations of the study

Because of the limited neuron and bouton data, the connectivity is very sparse on a whole-brain scale. Thus, the total number of putative detected boutons was 3,825,227, of which 181,691 (4.7%) were identified as potential connections with dendrites in our dataset.

There is a bias in the number of neurons on cell type. Only three types of neurons, VPM (385), CP (325), and SSp (253), had numbers above 100 and these three cell types accounted for one-half of the total data (1891). This leads to a specific description of the thalamo-cortical-striatal circuit.

The great variation in the size of the original brain leads to some stretching and shifting of the neuron in the registration. Some of the neurons, especially the neurons near the surface of the cortex, had some axons beyond the atlas volume. According to our statistics, there are 738 neurons with data points beyond the CCFv3 boundary. Among these neurons, the number of out-of-bounds points is 7.46% of all points. There is no better registration solution currently unless mass manual proofreading is used. Still, soma locations all lay within the atlas volume, and our connectivity generation method only requires axons and dendrites colocalization regardless of their location in space.

Due to data limitations, we can only get the bouton location on the presynaptic axon and do not have information about the spine on the postsynaptic dendrite.

## Acknowledgments

We thank Zhixi Yun, Feng Xiong and Lijun Wang for comments on the figures. This work was mainly supported by a Southeast University (SEU) initiative of neuroscience awarded to H.P.. H.P. was also supported by a Zhejiang Lab BioBit Program visiting grant (2022BCF07).

## Author contributions

H.P. and L.MG. conceived and designed the study. S.J. generated and provided the full morphology bouton data. P.Q. and L.MG. achieved all analysis results and wrote the paper with help from all authors.

## STAR★Methods

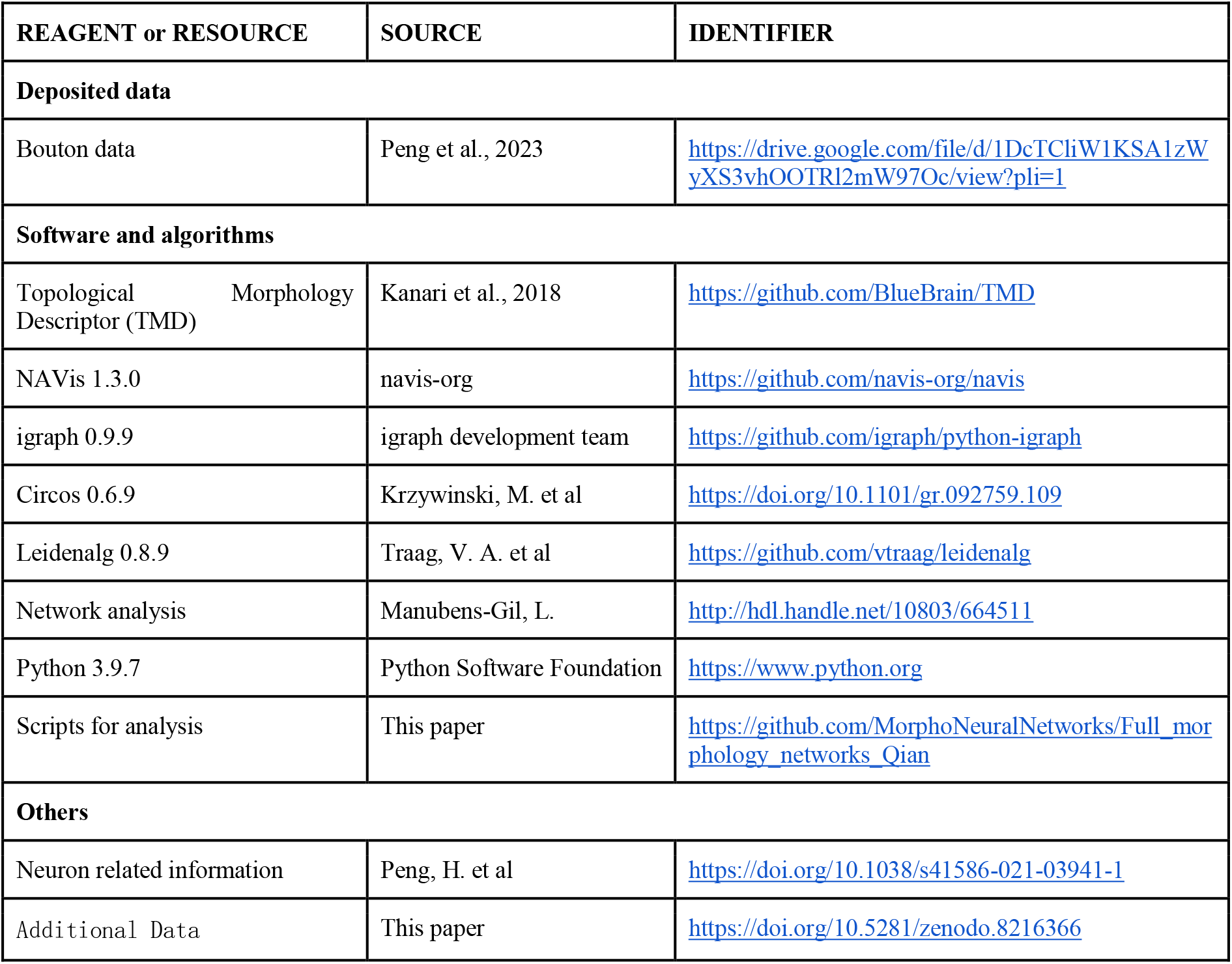
Key resources table.

## Resource availability

### Lead Contact

Further information and requests for resources and code should be directed to and will be fulfilled by the lead contact, Linus Manubens-Gil.

### Materials availability

This study did not generate new unique reagents.

### Data and code availability

All original data including full neuron reconstructions, together with observed bouton locations can be found at: https://drive.google.com/file/d/1DcTCliW1KSA1zWyXS3vhOOTRl2mW97Oc/view?pli=1.

The additional data like average bouton density, generated connectivity matrices,pertubation results can be found at https://doi.org/10.5281/zenodo.8216366.

All original code is publicly available at GitHub, https://github.com/MorphoNeuralNetworks/Full_morphology_networks_Qian.

### Method details

#### Sources of experimental data

Recent advances in light microscopy allowed the generation of complete neuronal reconstructions at micrometric resolution. Here, we used 1891 full neuron reconstructions data with axonal bouton locations from a dataset obtained at SEU-Allen (Peng et al., 2021). The data was generated using the MorphoHub platform (Jiang et al., 2022; Peng et al., 2023), which follows a multi-level annotation protocol that we describe briefly: First, the neuronal reconstruction is delineated using Vaa3D Terafly (Bria et al., 2016) and TeraVR (Wang et al., 2019), being cross-checked by at least two annotators. Axonal bouton data was obtained as described in (Liu et al., 2023). Finally, we remove any possible duplicates by deleting boutons at a distance closer than 5 voxels. Putative bouton locations are stored as an extra column in extended Stockley-Wheal-Cole (ESWC) files describing the neuron morphology (Ascoli et al., 2023).

The SEU-ALLEN dataset has a total of 1891 neurons suitable for predicting bouton locations. The dataset includes 97 cell types defined by the brain region where their soma is located (s-types; see glossary in Supplementary Material). The reconstructions have been obtained from 39 brains and are equally distributed between the left and right hemispheres. After annotation of the full neuronal structure, the trees have been spatially registered to CCFv3 as described in Qu et al. (Qu et al., 2022).

#### Calculation of bouton distribution

Since downsampling was used to reduce the file size after the boutons were identified, we resampled the ESWC files at an interval of 10 µm. We measured the distribution of bouton and axon length through the full axonal tree using the Sholl analysis on the neuron reconstructions before registration to CCFv3. The Sholl Analysis is the process of measuring neuron properties in concentric circles around the soma, and it provides a quantitative description of morphological features for the analyzed neurons (Sholl, 1953). We measured the number of branches intersecting each circle and both cable length and number of axon boutons between consecutive circles at 100 μm intervals. To do so, we used the “sholl_analysis” function of the Navis package (version 1.3.0) (Costa et al., 2016) in Python (version 3.9.7).

To validate the observed densities of boutons and to be able to compare to experimental measurements obtained with electron microscopy, we obtained average bouton densities (number of boutons per micrometer of axon length) for all axons located in specific CCFv3 regions. We obtained the total number of observed boutons and divided by the total axon length in each region.

To test the impact of the inhomogeneous observed bouton distributions on the network structure, we generated model neurons with homogeneous bouton densities as would be expected from using electron microscopy data to generate the connectivity. Given that different s-types showed diverse observed bouton distributions, we obtained the average bouton density for each s-type. To do so, we scaled the average axon length distribution of each s-type within a scaling factor representing bouton density in a range between 0 and 1 with steps of 0.001. 0 would imply no boutons at all throughout the tree, and 1 would imply one bouton for every micron of axonal length. We chose the scaling factor value that minimized the squared difference to the observed bouton distribution based on our experimental data. Given that these average bouton densities could be useful for generating connectivity in models of cortico-thalamic circuits the obtained values can be found in the Supplementary Material.

To simulate a uniform distribution of axonal boutons in the individual reconstructions according to the average density of each s-type, we devised an algorithm to define axon bouton locations synthetically. Specifically, first, we found all end nodes of all branches, which are leaf nodes and backtracked from these leaf nodes sequentially. In the process of backtracking, boutons were set at equal intervals defined by the inter-bouton distance determined by the inverse of the average bouton density. To prevent repeated assignment of boutons in low-order branches, paths that had already been traversed were not visited again.

In order to compare with the bouton density in previous experimental studies, we counted the total axon length and the bouton number from neurons with specific soma regions within any brain region in the CCFv3 model. The number of boutons per unit distance was obtained by dividing the two values. But such a result tends to underestimate the bouton density because boutons are not evenly distributed over the axon.

#### Generation of networks

Given that our full neuron reconstructions and bouton data are spatially registered to the mouse CCFv3, the neuronal morphologies can be explored in the same coordinate space, allowing us to explore the colocalization of axonal boutons and dendritic trees. We developed an algorithm to obtain a whole brain connectivity matrix at the single-cell level based on our dataset. In the resulting network, nodes are single neurons and edges are the connection strength between a pair of neurons *i* and *j*. According to Peters’ rule (Peters and Feldman, 1976), whether two neurons are connected can be determined by the presence of a nearby axon and dendrite. Here, we used a nuanced Peter’s rule (Rees et al., 2017) given that the potential connectivity is weighted by the number of boutons on such an axon-dendrite connection pair.

Specifically, first, we divided the whole brain into 30*30 μm cubes and calculated the axon length, dendrite length, and bouton number of each neuron within each cube. We considered that when both axons and dendrites are present in a cube with existing boutons, there is a potential connection. We defined the connection strength based on the number of boutons in each cube (Equation 1). Given that multiple pre-and postsynaptic neuron segments may coexist in each cube, we distributed the total number of observed boutons to all dendrites in the cube according to the proportion of dendrite length contributed by each neuron. This provides all pairwise connection strengths between pre-and postsynaptic neurons in each cube. By iterating this process through the whole brain, a full single-cell connectivity matrix is obtained.

The three networks used for comparison: the random network, the small-world network and the scale-free network, can be generated directly by the functions in Igraph: “Erdos_Renyi()” (Erdős and Rényi, 1960), “Watts_Strogatz()” (Watts and Strogatz, 1998), “Barabasi()” (Barabási and Albert, 1999). For the random network, we keep the number of nodes and edges the same as for the *observed* network. For the small-world network, we set the dimension of the lattice to 1 and choose the rewiring probability to be 0.02. The size is the number of nodes in *observed* network. And the number of edges is adjusted by giving the distance (number of steps) within which two vertices will be connected to make that as close as possible to the *observed* network. Finally the extra edges are removed randomly. Similarly, for the scale-free network, we adjust the number of outgoing edges generated for each vertex to approximate the *observed* network while keeping the number of nodes the same, and finally remove the excess edges randomly as well.

#### Network analysis

To quantify network structural properties, we obtained the degree distribution, triad census, hubs, and authorities using the “igraph” package (version 0.9.9) in Python (version 3.9.7). Correspondingly, this toolkit provides these functions: “degree_distribution()”, “triad_census()”, “authority_score()”, “hub_score()”, which we used with default parameters. We generated circular plots to visualize the networks using Circos (Krzywinski et al., 2009) (version 0.6.9).

#### Community detection

To explore the community structure of the networks, we used the Leiden algorithm (Traag et al., 2019), which is an optimization of Louvain’s clustering method (Blondel et al., 2008) that ensures detected communities are connected and have faster computation. Specifically, the Leiden algorithm divides the graph nodes into communities while optimizing modularity in three phases: (1) local assignment of nodes into communities, (2) refinement of the partition, and (3) aggregation of the network reducing the number of nodes to represent communities. Here we used the “leidenalg” package (Traag et al., 2019) (version 0.8.9) in Python (version 3.9.7), and to ensure that the analyzed communities had a sufficient number of nodes, we arbitrarily selected the largest six groups for subsequent analysis and especially to generate the simple plots accounting for the major communities in each network.

#### Multi-objective optimality metrics

Considering that the basic function of a neural network is the transmission and storage of information and that building a network has a material and metabolic cost, we use three quantities to explore putative functional constraints of the network: cost, storage capacity, and routing efficiency (Manubens-Gil and others, 2018).

#### Cost

Cost is defined as the number of boutons in the network.

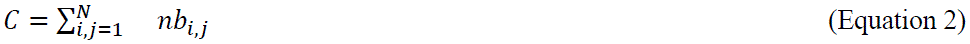

where *nb_i,j_* is the number of boutons connecting a pair of neurons *i* and *j*, and *N* is the total number of neurons in the network.

#### Storage Capacity

We estimated the storage capacity of a network as the sum of the total number of non-redundant possible states for each neuron receiving *s* connections provided by *d* pre-synaptic neurons as previously defined by Poirazi and Mel for linear neurons (Poirazi and Mel, 2001). Briefly, the combinatorial “n choose k” quantification of possible states for a post-synaptic neuron expressed in bits (basic unit of information) is given by:

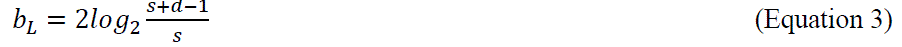

and total storage capacity of the network by:

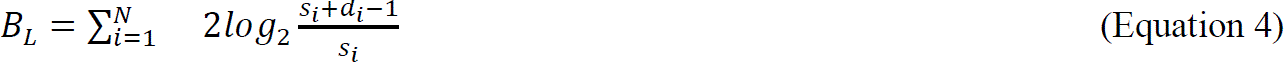

Where *N* is the number of nodes in the network.

#### Routing Efficiency

Routing efficiency is inversely proportional to the weighted shortest path length φ_*ij*_in the network between any pair of nodes (neurons) *i* and *j*. When two neurons in the network are closely connected or have more synapses, the path length between them is reduced, and the routing efficiency increases. We obtained the shortest path length matrices using an in-house implementation of the Floyd-Warshall algorithm (Floyd, 1962; Roy, 1959; Warshall, 1962). The formal definition for the routing efficiency is as follows:

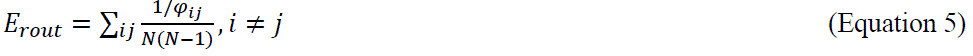

Where φ*ij* is the graph shortest path length between the nodes *i* and *j*, and *N* is the total number of nodes in the graph.

#### Perturbation

To study the effect of putative biologically realistic (Kulkarni and Firestein, 2012) morphological perturbations on the network, we designed three perturbation operations: scaling of neuron size, pruning of neuron branches, and removal of synaptic boutons. We used those to perturb morphological details of the neurons, including their size, complexity of the neurites, and number of boutons.

#### Scaling of neuron size

This operation involves reducing the size of the axonal, dendritic tree, or entire neuron by a factor ranging from 0.5 to 0.9 in intervals of 0.1. For dendrite scaling, we select dendritic branches and scale the 3D spatial coordinates of all points forming the branches relative to the coordinates of the soma. Since all dendritic branches in our data are connected to the soma, we can accurately scale their coordinates. For axon scaling, we identify the longest axon branch as the projection branch and keep it unmodified. Then, we scale the coordinates of the axon subtrees relative to the point of connection to the projection branch.

To separately study the impact of bouton distribution and axonal tree complexity, we adjust the number of boutons when scaling axons. Bouton locations are assigned to specific nodes in the neuron tracings. In the case of uniform distribution, we reset the position of the boutons in the scaled axon according to the bouton density per unit of axon length. In the case of observed boutons, we first calculate the distance between each consecutive pair of boutons and sort them from smallest to largest. The number of boutons to be deleted is determined based on the scaling ratio, and the boutons are uniformly deleted from the distance-ordered list. This procedure ensures that the distribution of boutons per unit of length remains unchanged after scaling the axonal tree size.

#### Pruning of neurites

Neurite pruning refers to the process of deleting a certain percentage of dendritic or axonal branches to modify the neuron morphology. In our study, we performed various types of perturbations by removing only the axonal, dendritic branches, or both. First, we identified the number of leaf nodes (termination points) in a given neuron. We then selected a set of leaf nodes based on a pruning ratio ranging from 0.5 to 0.9 in 0.1 intervals. The pruning of single branches began from one of the selected leaf nodes and proceeded through parent nodes until the first branch node was encountered, and all nodes in the path were deleted.

#### Removal of boutons

The process of removing boutons randomly does not alter the neuron morphology but only deletes a fixed percentage of boutons. We start by identifying the total number of boutons in a neuron, after which we select boutons randomly at a fixed proportion range between 0.5 and 0.9 in ten percent intervals. We label the selected points as axonal continuation points, which signifies that the reassigned nodes are not considered while generating connectivity matrices to establish connections.

## Supplementary Materials

**Table S1**

Full names of all cell types involved, acronyms, number of neurons, and average bouton density.

**Table S2**

Comparison between the bouton density calculated from our data and other articles.

**Table S3**

Statistical tests in Figure 3, including correlation and independence statistical test of degree distribution and triad census among different networks.

**Table S4**

Network analysis results: average path length+clustering coefficient; hubs and authorities scores and triad census.

**Supplementary Figure 1.**
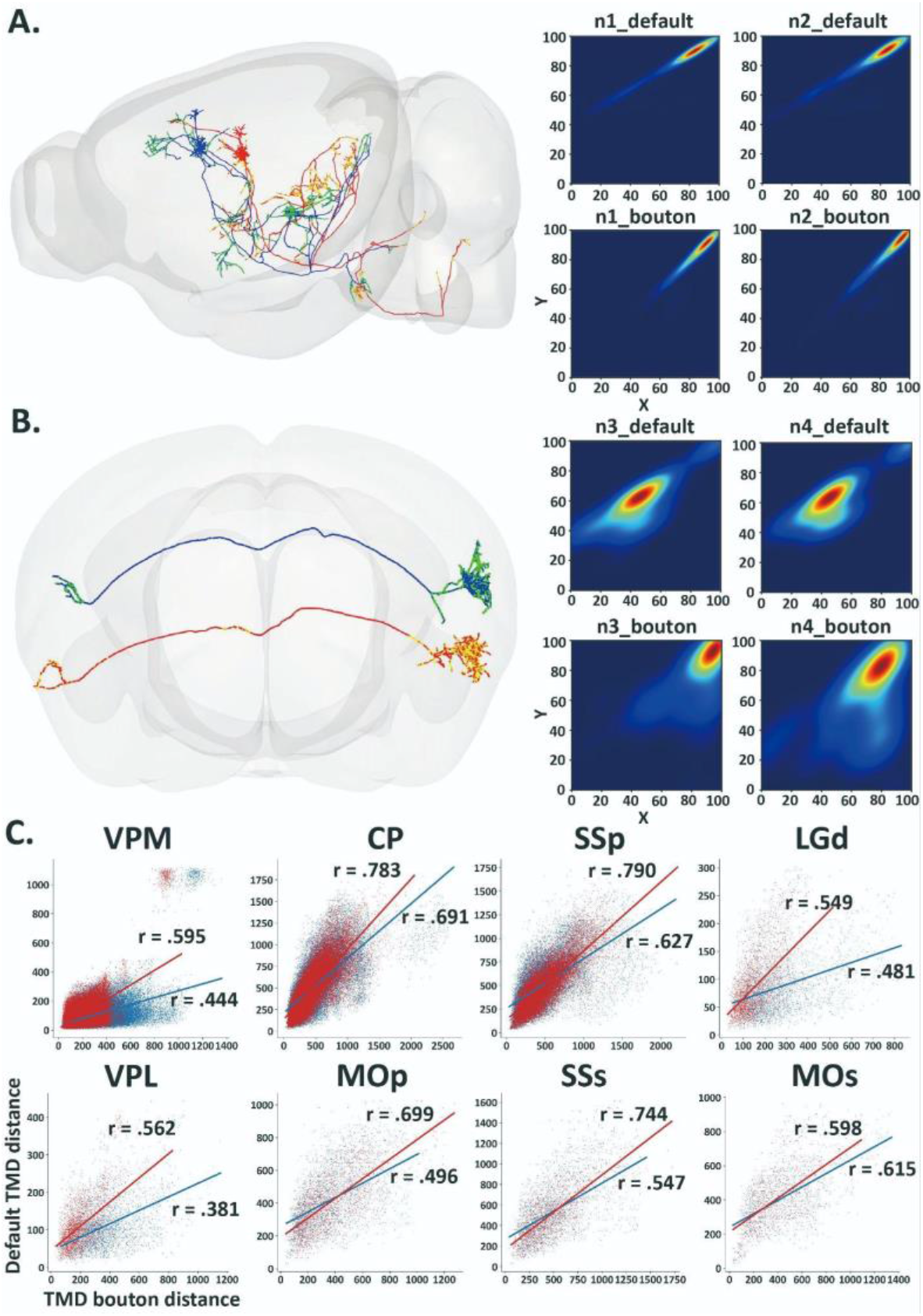
Related to Fig. 1: TMD results for other cell types and two examples. **(A, B)** One pair of neurons (red vs blue) with similar morphology and bouton distribution (yellow vs green, top) and one pair of neurons with similar morphology but non-similar bouton distribution (bottom). On the left is the persistence diagram for axon and bouton. **(C)** Scatterplot of the Topological Morphology Descriptor (TMD) distances between pairs of neurons with somas in the same region. We calculated distances by defining the persistence histogram as the number of boutons within the radius (TMD bouton; x axis) or the distance to soma (TMD default; y axis) at birth and death points of neuron segments. The plot shows pairwise distances for neurons with the measured (observed) bouton locations (in blue) and analogous measures for uniform distributions of boutons randomly located throughout the axonal tree (in red). And this phenomenon exists in different cell types.

**Supplementary Figure 2.**
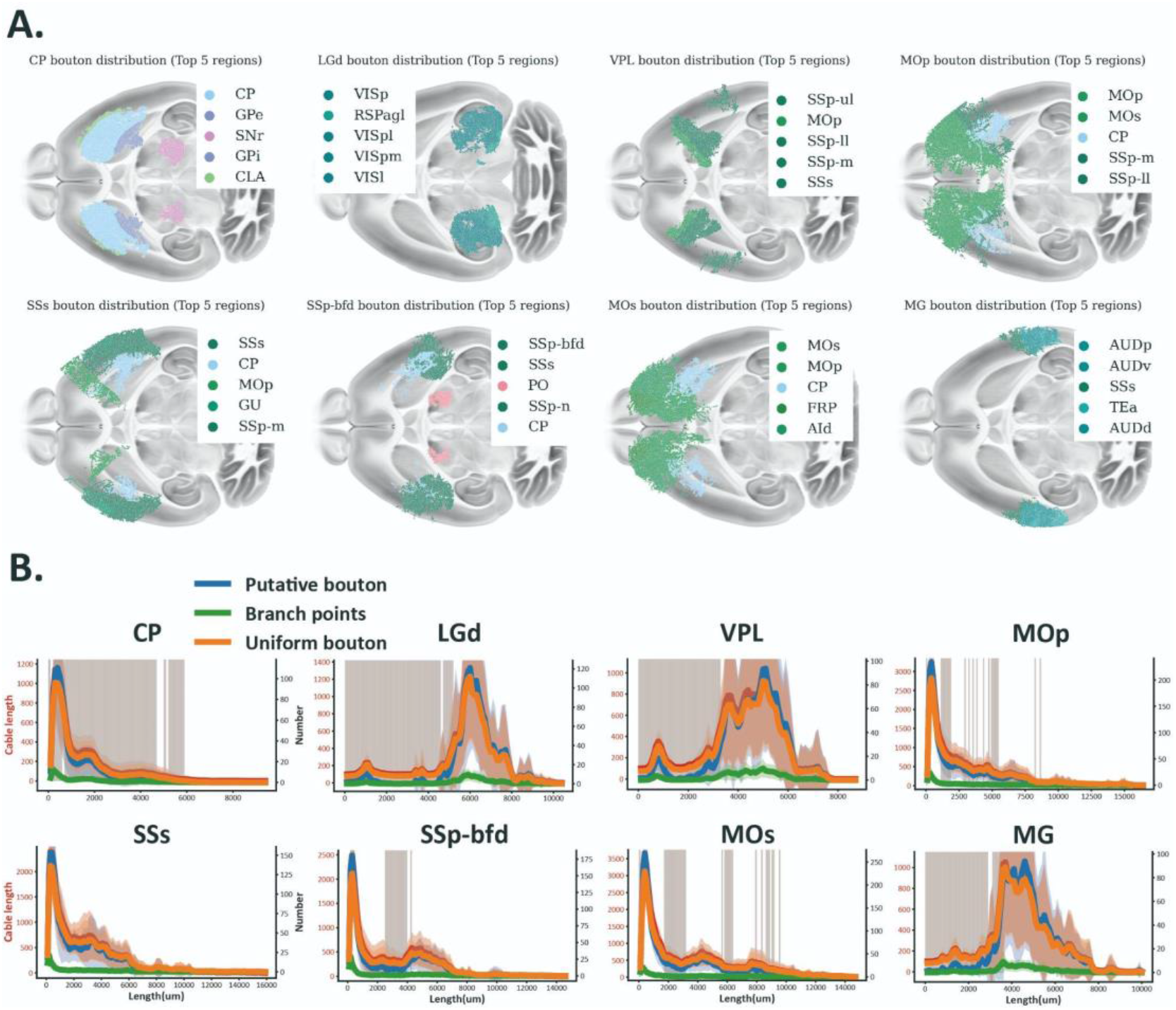
Related to Fig. 1: Visualization of bouton distribution and sholl analysis of other cell types. **(A)** 2D horizontal projection of bouton locations in the top five regions most innervated by different cell types. Cortical regions are colored in shades of green, caudoputamen (CP) in the Striatum, in blue, and VPM in the Thalamus in pink. **(B)** Sholl analysis of the difference between a uniform bouton distribution (blue) and the observed bouton distribution (orange) for neurons with their soma in different regions. Statistically significant differences between the two distributions are indicated with a grey shadow; paired-samples t-tests random vs. observed bouton number test p<0.005. Green lines indicate the number of branch points between Sholl concentric circles. Red lines show the Sholl analysis of axon length.

**Supplementary Figure 3.**
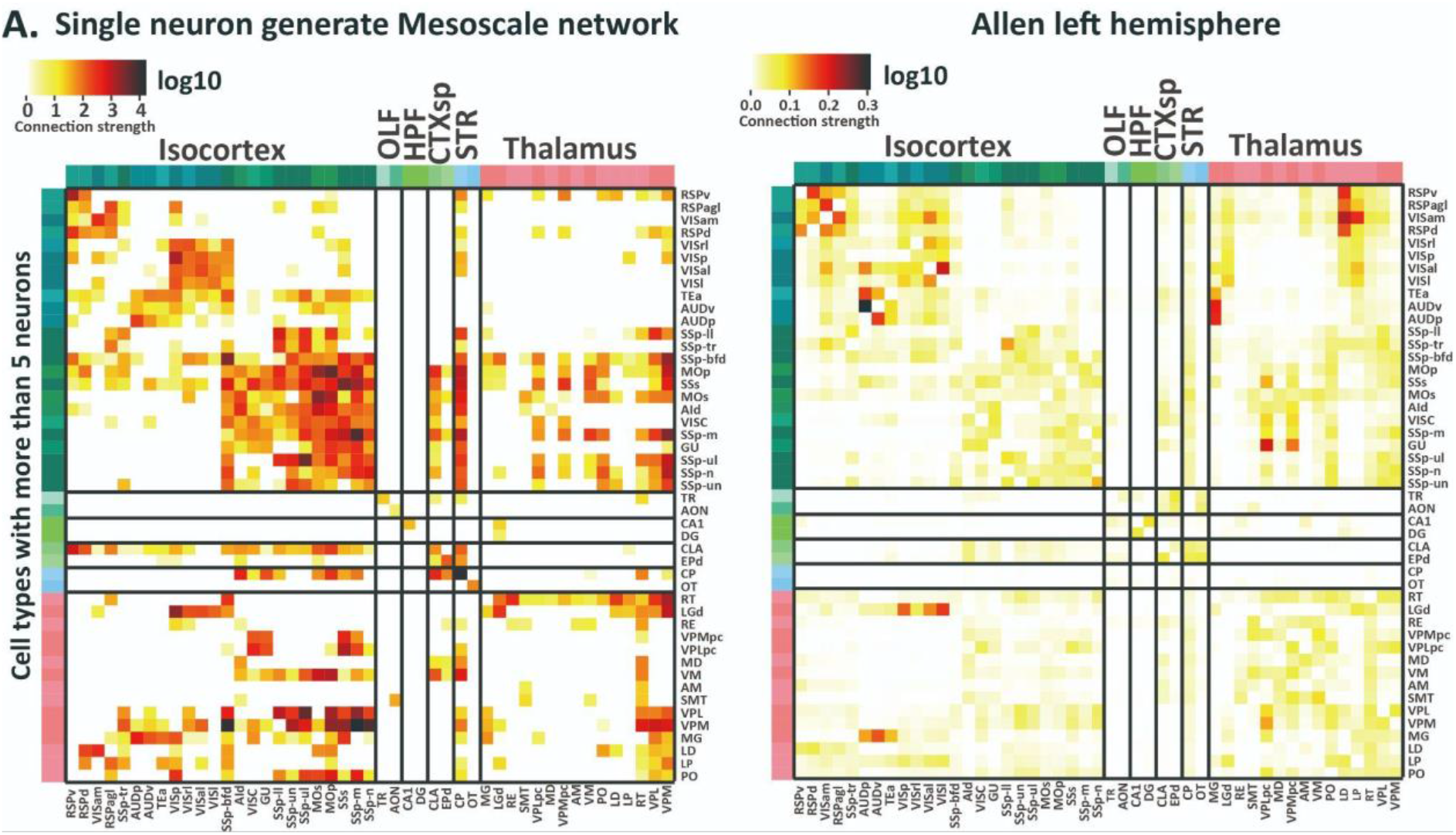
Related to Fig. 3: Comparison of mesoscale networks. (**A**) Heatmap of region connectivity consisting of single cell connections (left) and experimental mesoscale connections (right).

**Supplementary Figure 4.**
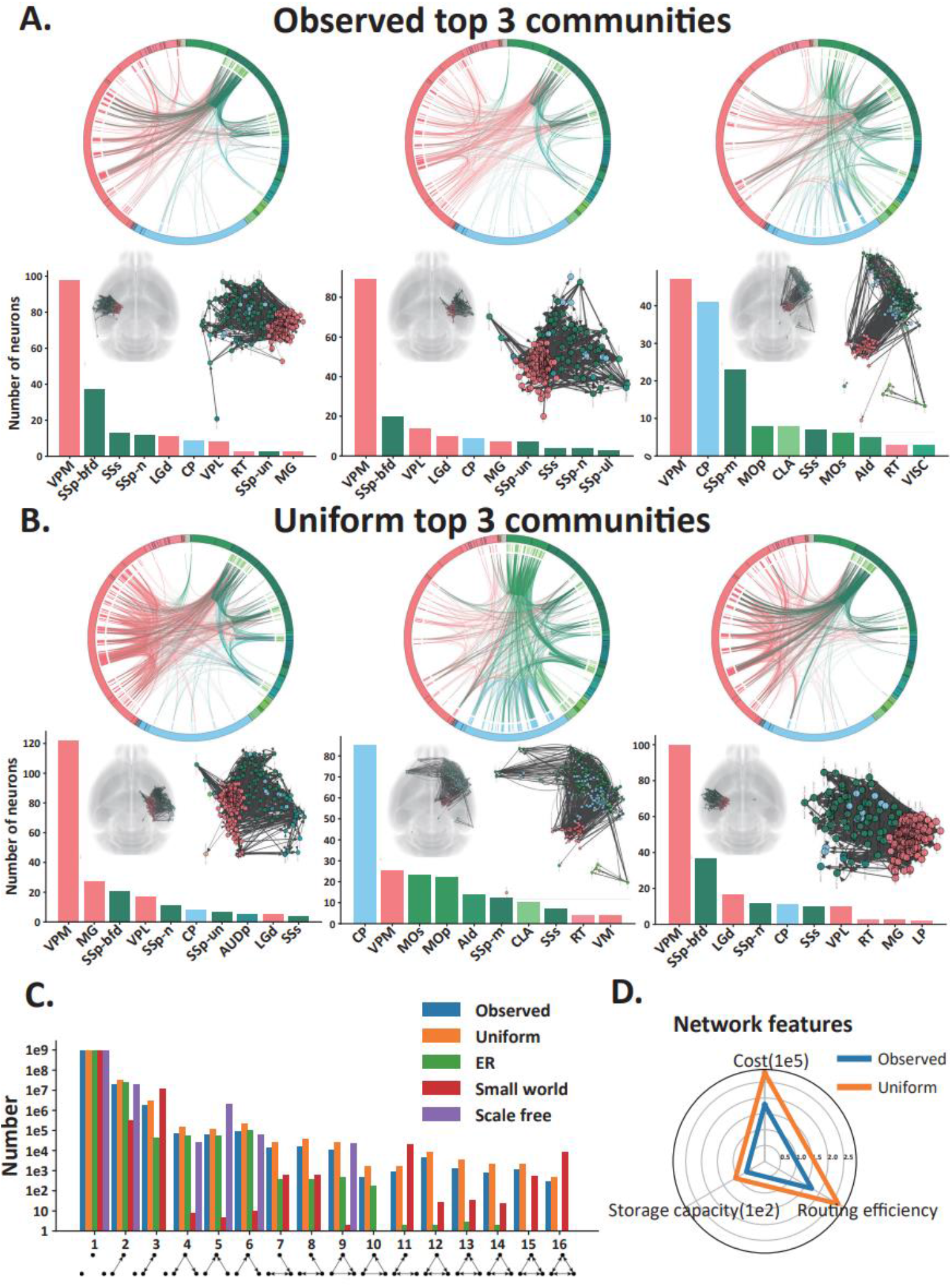
Related to Fig. 3: Supplementary results for *observed* and *uniform* network analysis. **(A, B)** Visualization of the first three communities of the *observed* and *uniform* networks. Cortical regions/connections are colored in green, caudoputamen (CP) in the Striatum, in blue, and VPM in the Thalamus in pink. The location of each community in the mouse brain is also shown. **(C)** Barplot of the number of triad occurrences with the same number of nodes and edges of *observed* (blue), *uniform* (orange), random (green), small-world (red), and scale-free (purple) networks. **(D)** Radar plot of the cost (number of potential connections), routing efficiency, and storage capacity of the *observed* (blue) and *uniform* (orange) networks.

**Supplementary Figure 5.**
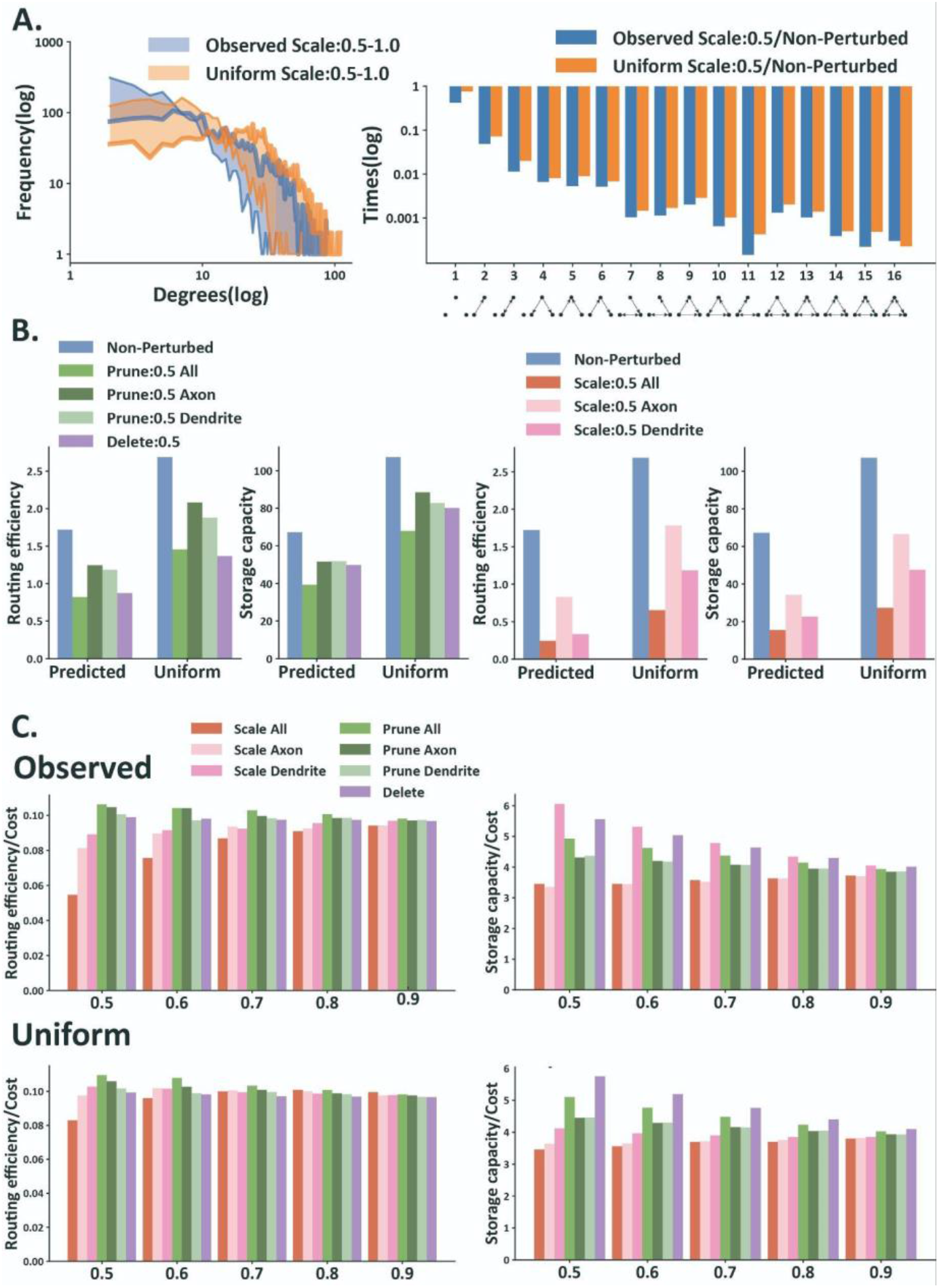
Related to Fig. 4: Additional results for perturbation network analysis. **(A)** The variation of the degree distribution and traid census of the *observed* (blue) and *uniform* (orange) network after scale. The shading is distributed at a scale varying from 0.5 to 1 at intervals of 0.1. **(B)** Bar plots of true value of routing efficiency (left) and storage capacity (right) for the unperturbed *observed* and *uniform* networks (blue) and after scale (red), prune (green) and bouton deletion (purple) perturbations. **(C)** Bar plots of routing efficiency (left) and storage capacity (right) divided by the cost (number of potential contacts) for the unperturbed *observed* and *uniform* networks (blue) and after scale (red), prune (green) and bouton deletion (purple) perturbations varying from 0.5 to 0.9 at intervals of 0.1. Bars indicate mean values ±standard deviation.

**Supplementary Figure 6.**
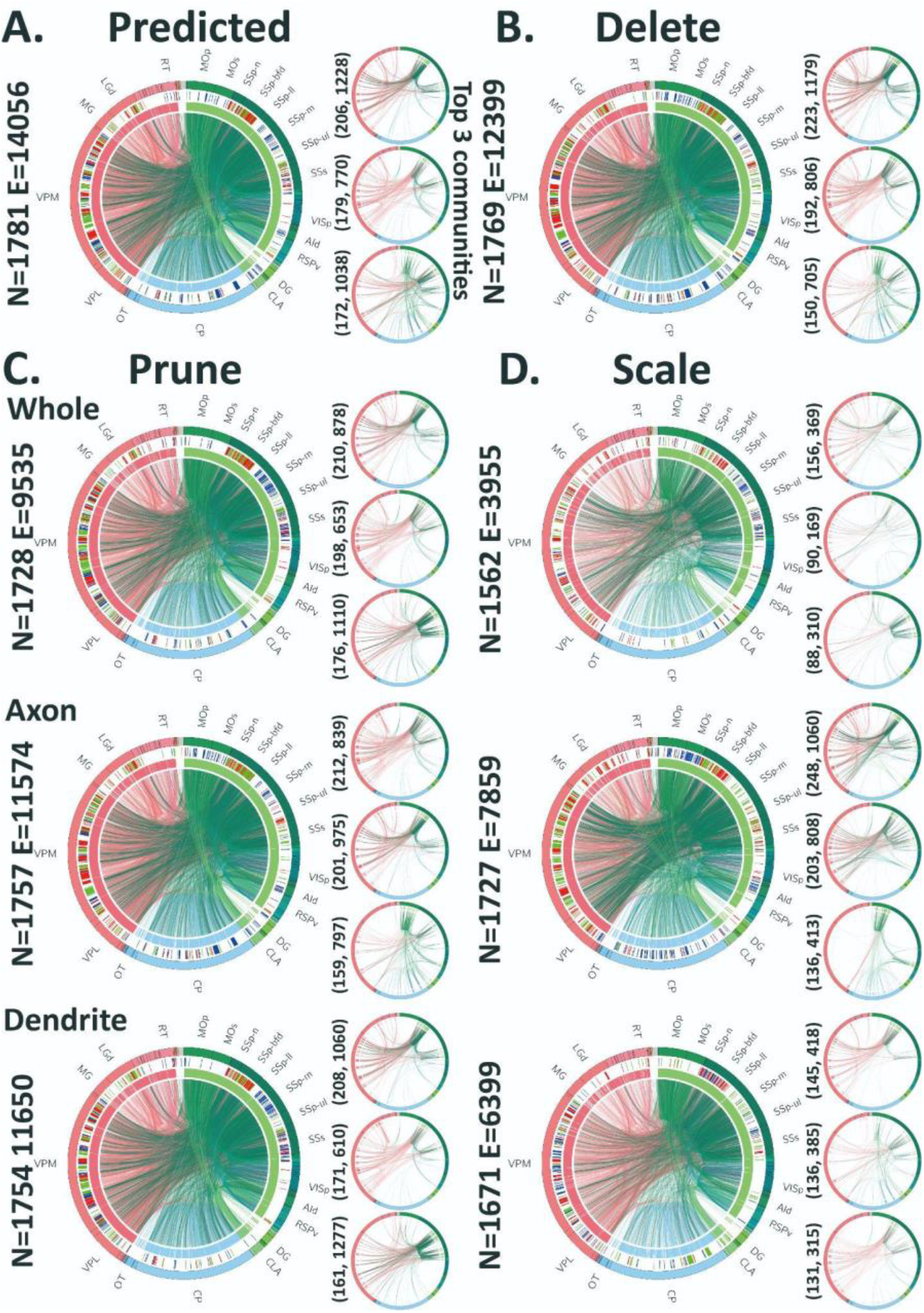
Related to Fig. 4: Community detection results for different perturbation network. **(A)** Circular plot visualization of the single-cell network based on the *observed* bouton data with its top 3 biggest communities. **(B, C, D)** Circular plot visualization of the single-cell network based on bouton data after three perturbations operations: scale, prune, delete bouton, which targeted on axon, dendrite, or complete neuron, with their top 3 biggest communities.

**Supplementary Figure 7.**
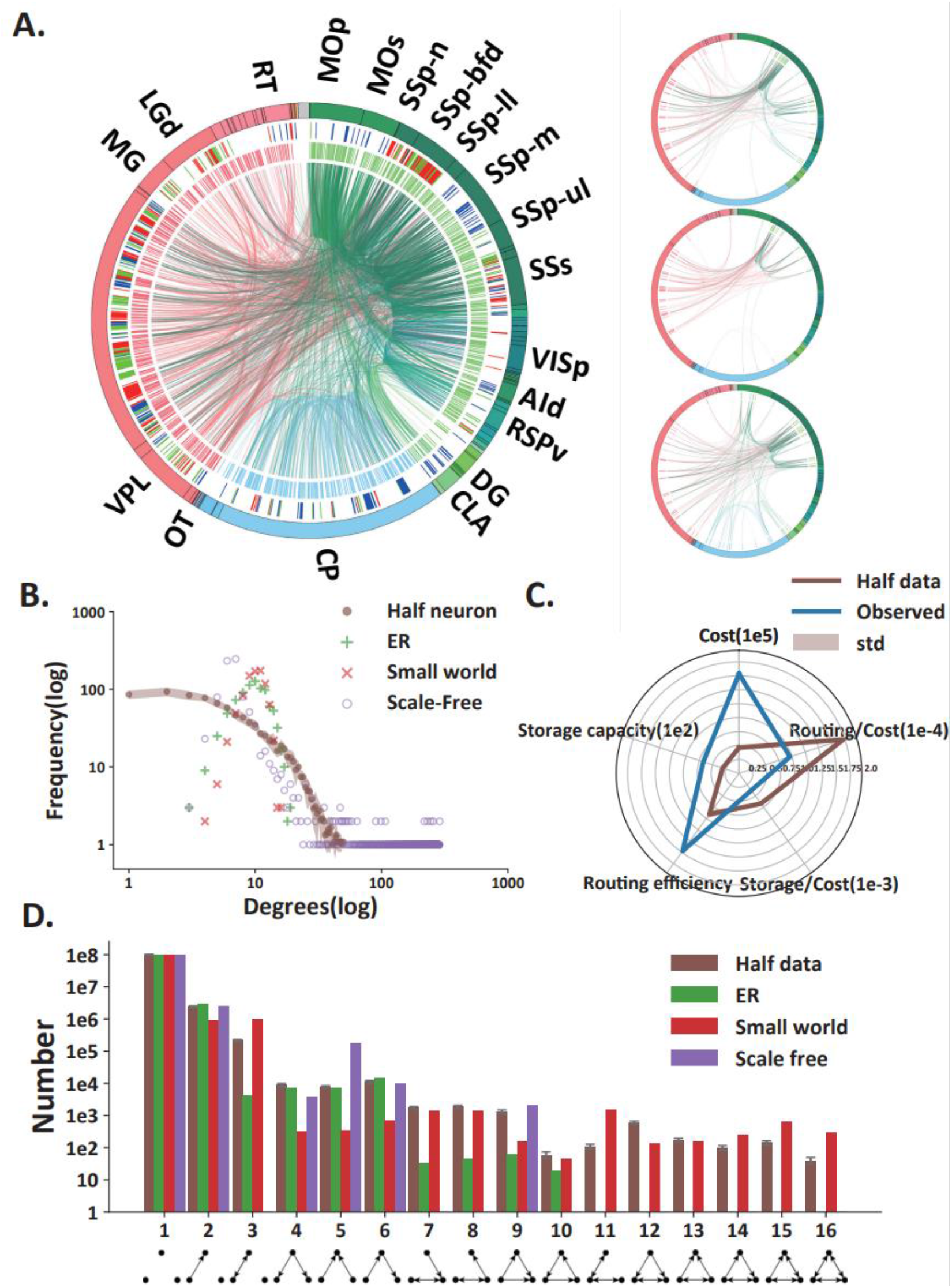
Related to Fig. 4: Results of constructing networks using half of the data. **(A)** Circular plot visualization of the single-cell networks constructed based on randomly selected half of neuron data (left). The left side is the top 3 communities in the network. **(B)** Results of network analysis, including degree distribution, triad census, cost, routing efficiency, and storage capacity, where the brown is the half data case. The trend is basically the same as the *observed* network.

