## Supplementary figures and images for "Non-homogenous axonal bouton distribution in whole-brain single cell neuronal networks"

### Supplementary Figure 1.pdf

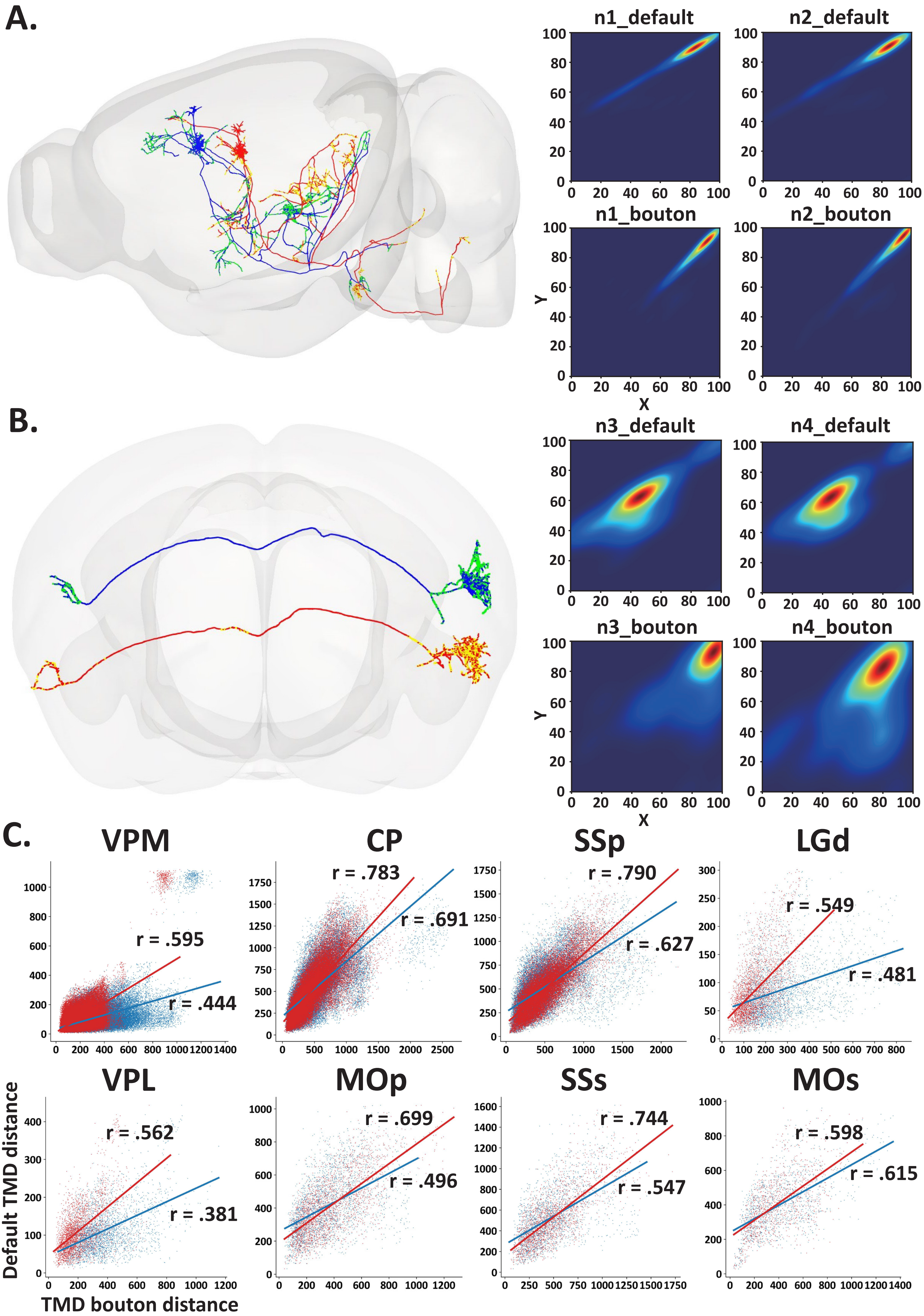

### Supplementary Figure 2.pdf

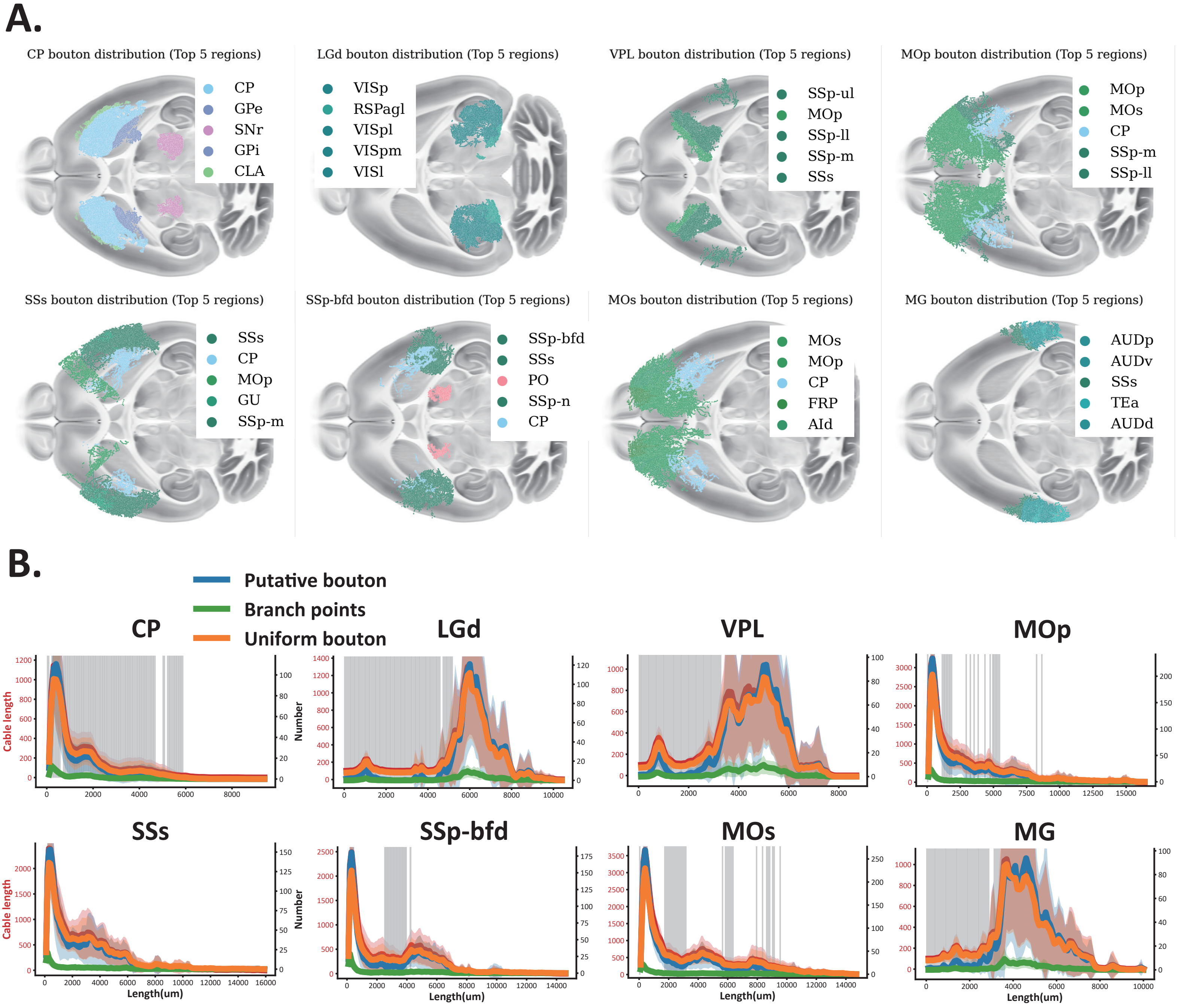

### Supplementary Figure 3.pdf

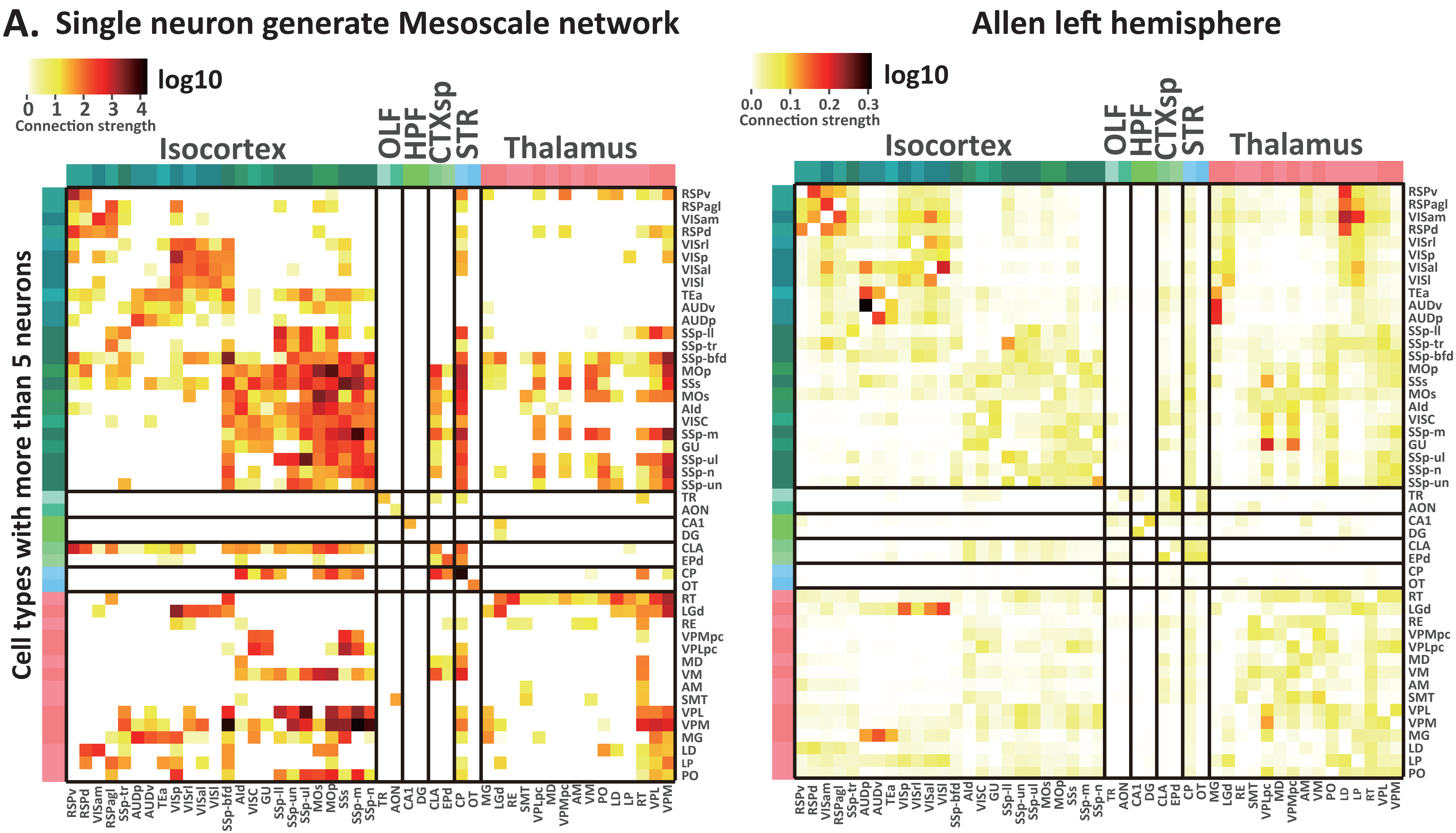

### Supplementary Figure 4.pdf

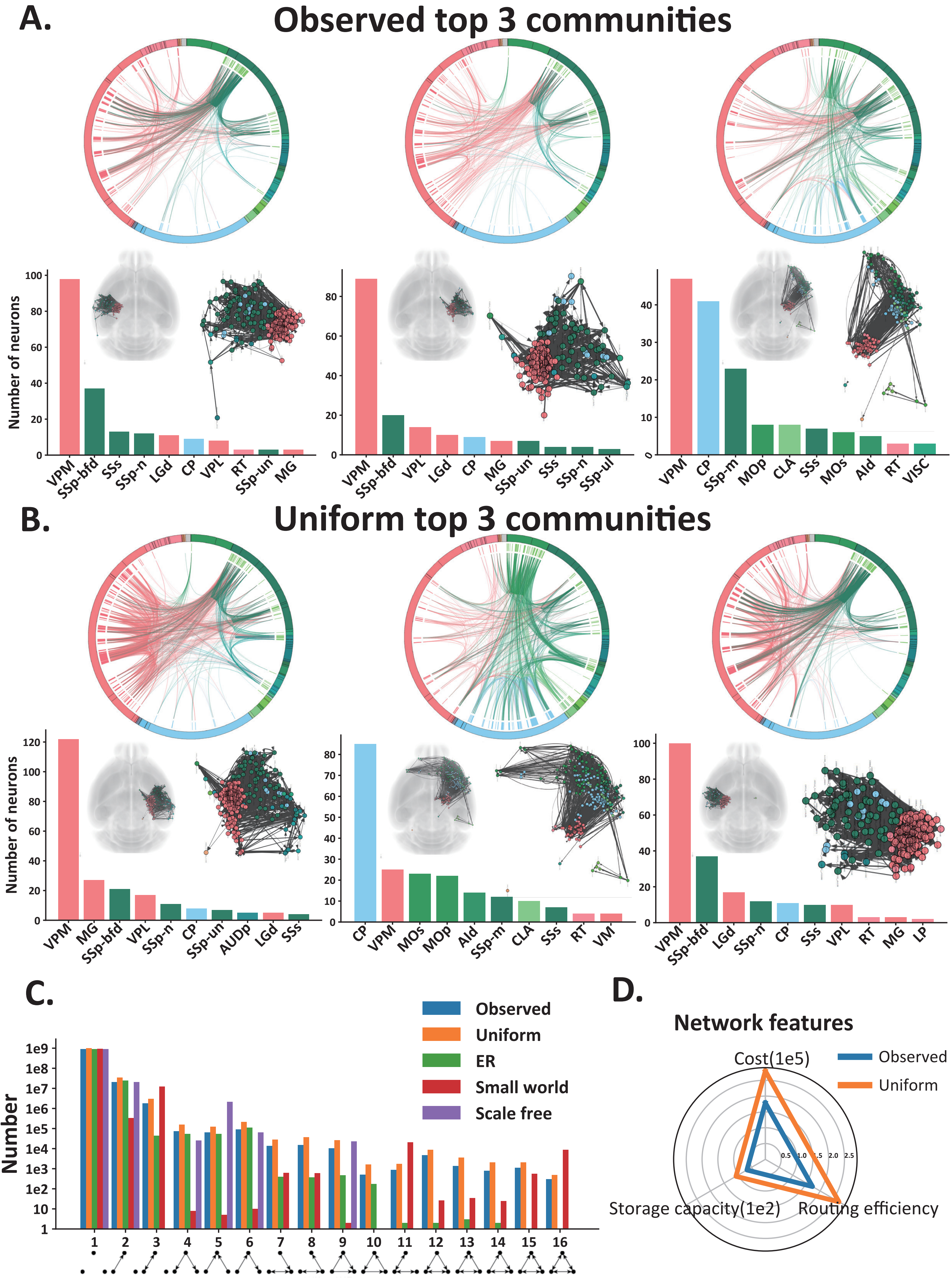

### Supplementary Figure 5.pdf

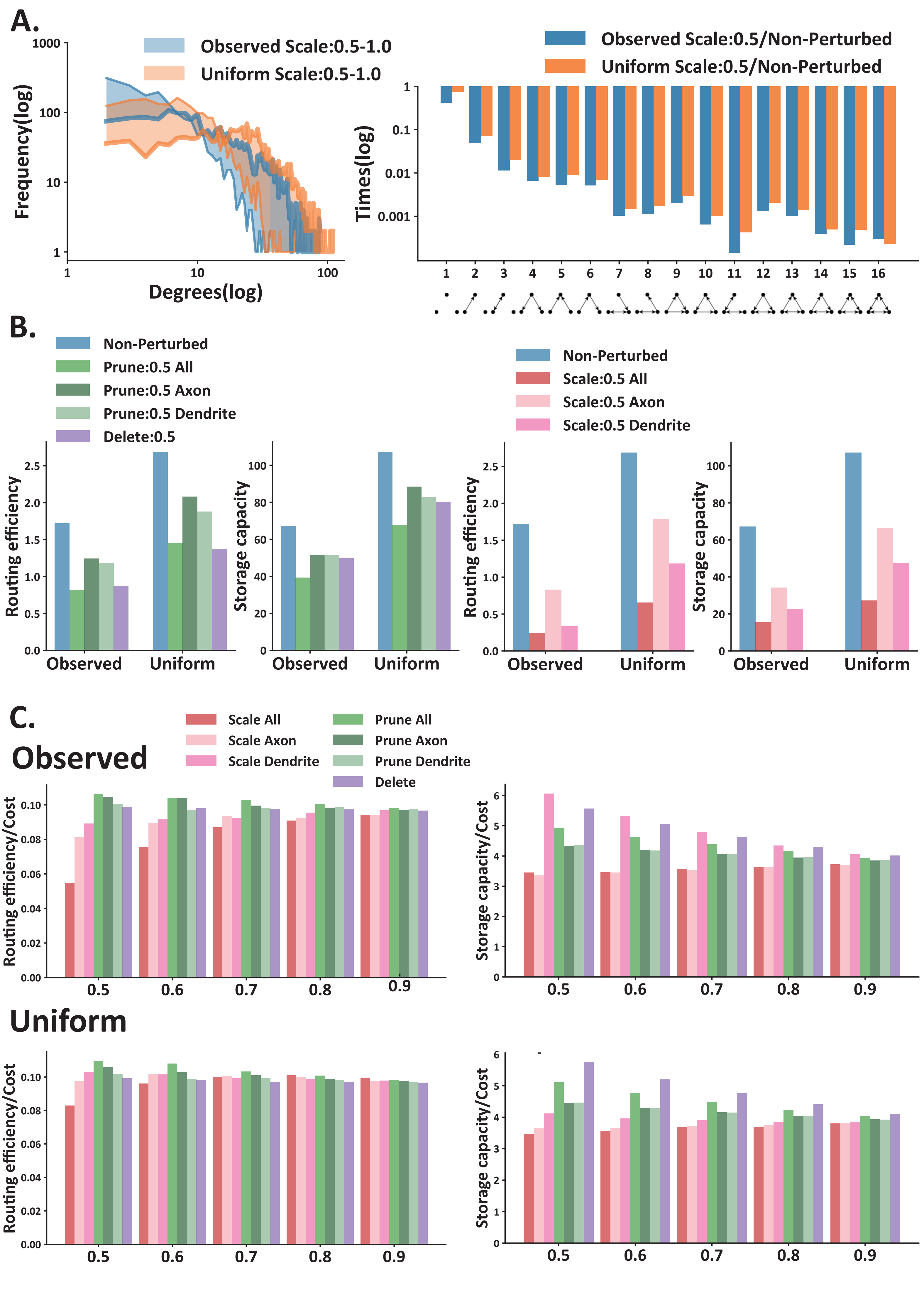

### Supplementary Figure 6.pdf

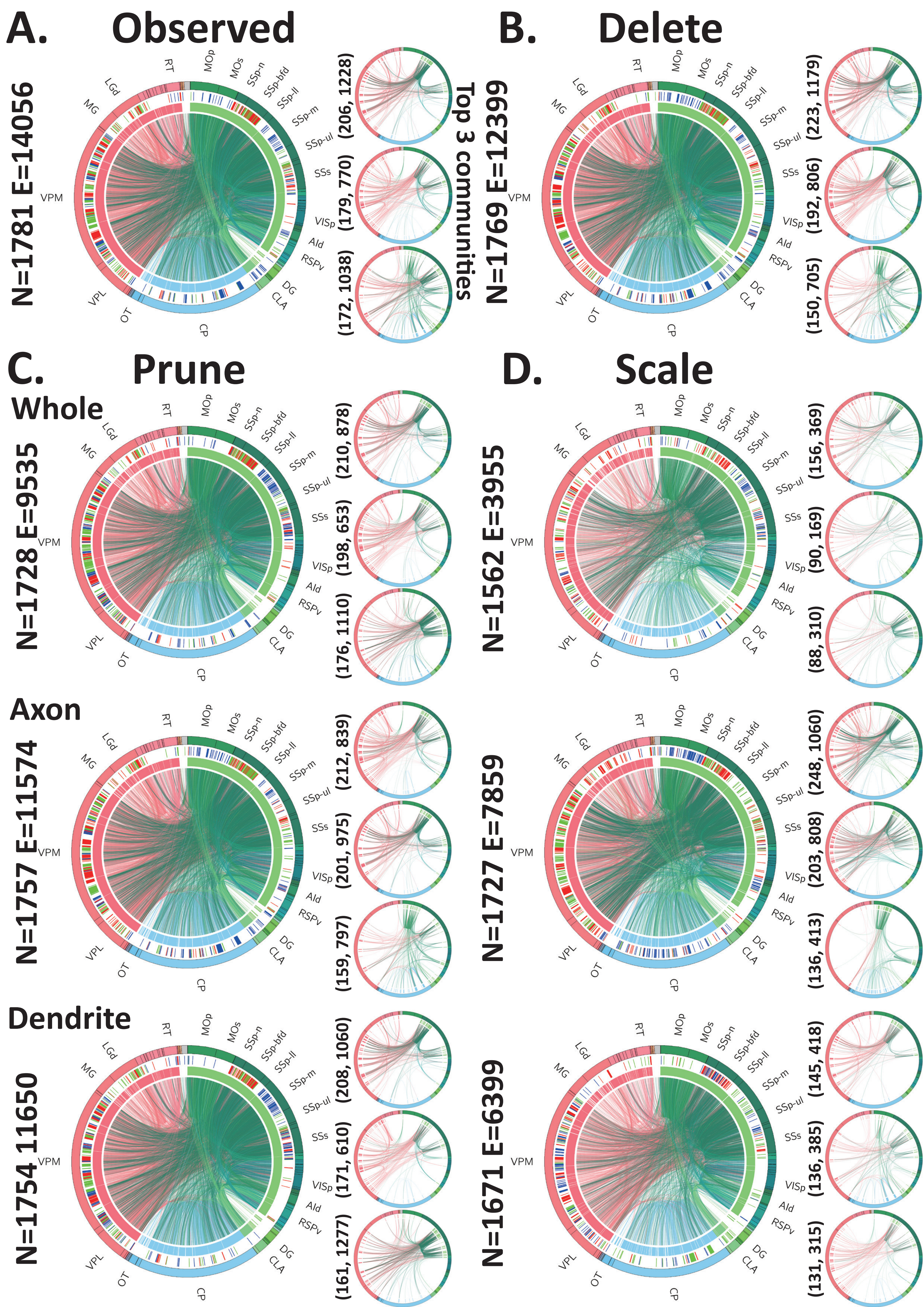

### Supplementary Figure 7.pdf

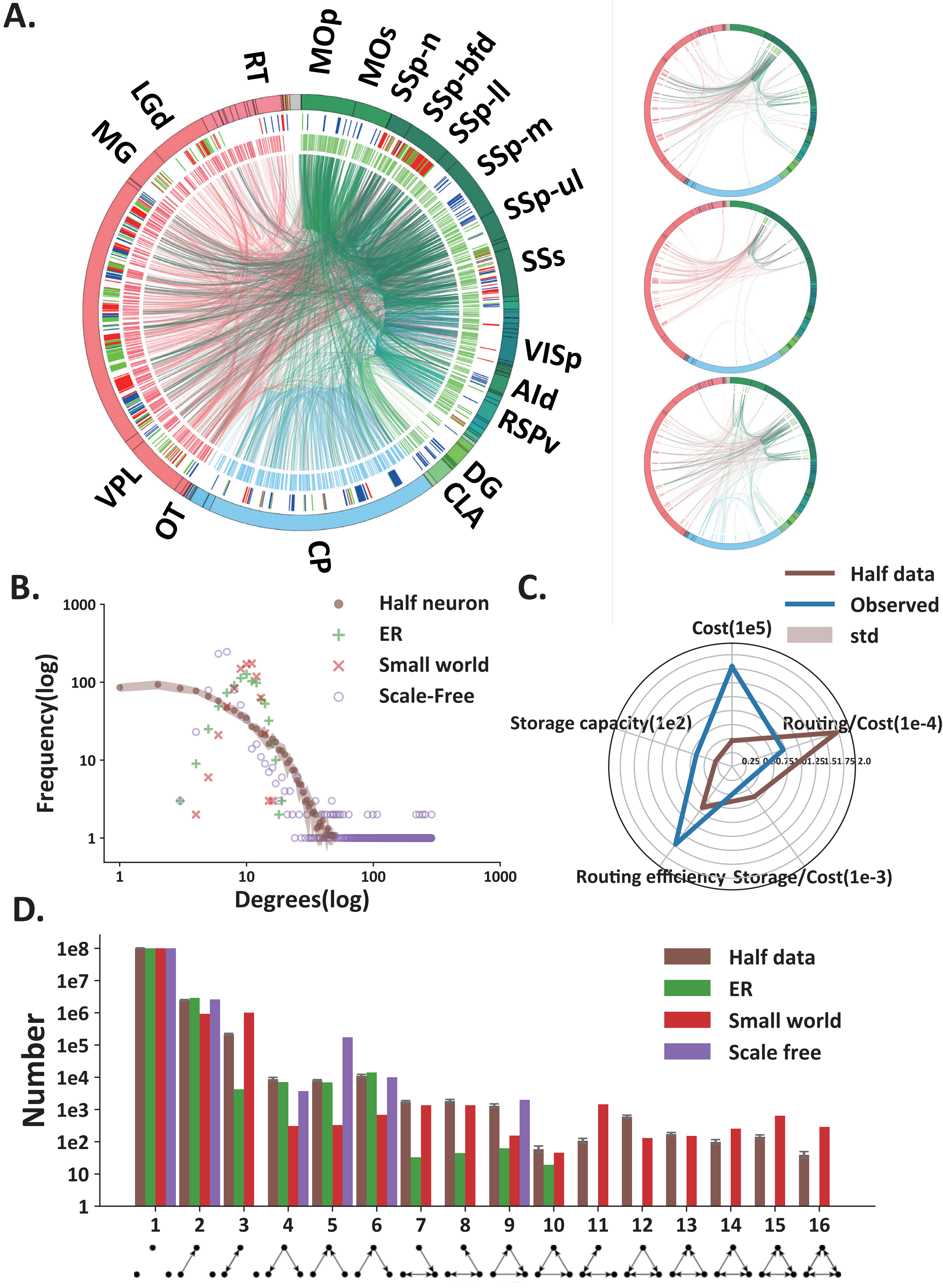
